# Vaccine antigen, Factor H binding protein, is typically a non-lipidated precursor that localises to the meningococcal surface by Slam

**DOI:** 10.1101/693374

**Authors:** RAG da Silva, AV Karlyshev, NJ Oldfield, KG Wooldridge, CD Bayliss, A Ryan, R Griffin

## Abstract

Meningococcal surface lipoprotein, Factor H binding protein (FHbp), is the sole antigen of the Trumenba vaccine (Pfizer) and one of four antigens of the Bexsero vaccine (GSK) targeting *Neisseria meningitidis* serogroup B isolates. Lipidation of FHbp is assumed to occur for all isolates and its surface localisation is conducted by surface lipoprotein assembly modulator, Slam.

We show in 91% of a collection of UK isolates (1742/1895) non-synonymous single nucleotide polymorphisms (SNPs) in the signal peptide of FHbp. A single SNP, common to all, alters a polar amino acid that abolishes processing, including lipidation and signal peptide cleavage. Rather than the toxic accumulation of the precursor in the periplasm as expected from disrupting the canonical processing pathway, remarkably the FHbp precursor is translocated to the outer membrane and surface-localised by Slam. Thus we show Slam is not lipoprotein-specific. In a panel of isolates expressing precursor FHbp at the surface, we investigated their binding to human factor H and their susceptibility to antibody-mediated killing. Our findings have implications for Trumenba and Bexsero and provide key insights for lipoprotein-based vaccines in development.

## Introduction

*Neisseria meningitidis* is a leading cause of bacterial meningitis and sepsis with high fatality (up to 50% when untreated) and high frequency (more than 10%) of severe sequelae [1]. Polysaccharide-based vaccines are effective in preventing disease caused by isolates of serogroups A, C, W and Y but are ineffective against serogroup B (MenB) isolates [2]. The lipoprotein, Factor H binding protein (FHbp), is a major virulence factor, which recruits human factor H (fH) to the meningococcal surface preventing complement from binding to the bacterium and thus inhibiting bacteriolysis by the alternative complement pathway [3].The amino acid sequence of FHbp varies with identities as low as 60% between isolates which led to the need for classification of this lipoprotein. FHbp was classified into subfamily A (subdivided into variant groups 2 and 3) and subfamily B (variant group 1) [4, 5] [6, 7]). Despite this variation, FHbp emerged as a promising vaccine candidate due to its ability to stimulate a strong serum bactericidal antibody (SBA) response capable of killing diverse group B isolates [4]. It is thought that FHbp-specific antibodies not only promote bactericidal killing by the classical pathway but also via amplification of the alternative pathway, by preventing fH from binding to FHbp [8].

Lipoproteins, such as FHbp, are synthesized as precursors (preprolipoproteins) in the cytoplasm, which are subsequently taken through a sequential pathway for processing and sorting to the outer membrane (OM) [9, 10]. The N-terminal signal peptide (SP), characteristic of bacterial lipoproteins, comprises a positively charged n-region, a hydrophobic h-region and a c-region with the consensus sequence [LVI][ASTVI][GAS] followed by an invariant C residue, known as the lipobox [11]. Translocation of the preprolipoprotein across the inner membrane (IM) occurs predominantly via the general secretory or Sec pathway [12]. Both the n-region and h-region are involved in interaction with SecA or other chaperones which deliver the precursor protein to the Sec-YEG transmembrane channel [13]. Preprolipoprotein diacylglyceryl transferase, Lgt, transfers the diacylglyceryl group from phosphatidylglycerol of peptidoglycan to the conserved C residue [14]. This diacylglyceryl modification of preprolipoprotein is vital for substrate recognition by the dedicated lipoprotein signal peptidase LspA [15] which cleaves the SP [16]. In diderms, such as Neisseria, the amino group of the C residue is further modified with a third acyl chain through apolipoprotein N-acyltransferase, Lnt [9]. Fully processed lipoproteins (cleaved and lipidated) that are destined to be anchored to the OM are transported by the Lol (lipoprotein outer membrane localization) machinery [17]. Lol consists of the IM ABC transporter-like complex, LolFD in *N. meningitidis* [9], the periplasmic LolA chaperone and the OM LolB lipoprotein receptor (Fig 1). FHbp is then surface localised by lipoprotein assembly modulator (Slam) by delivery to another OM protein or complex such as Bam or by acting as a conduit [18] (Fig 1).

**Figure 1.**
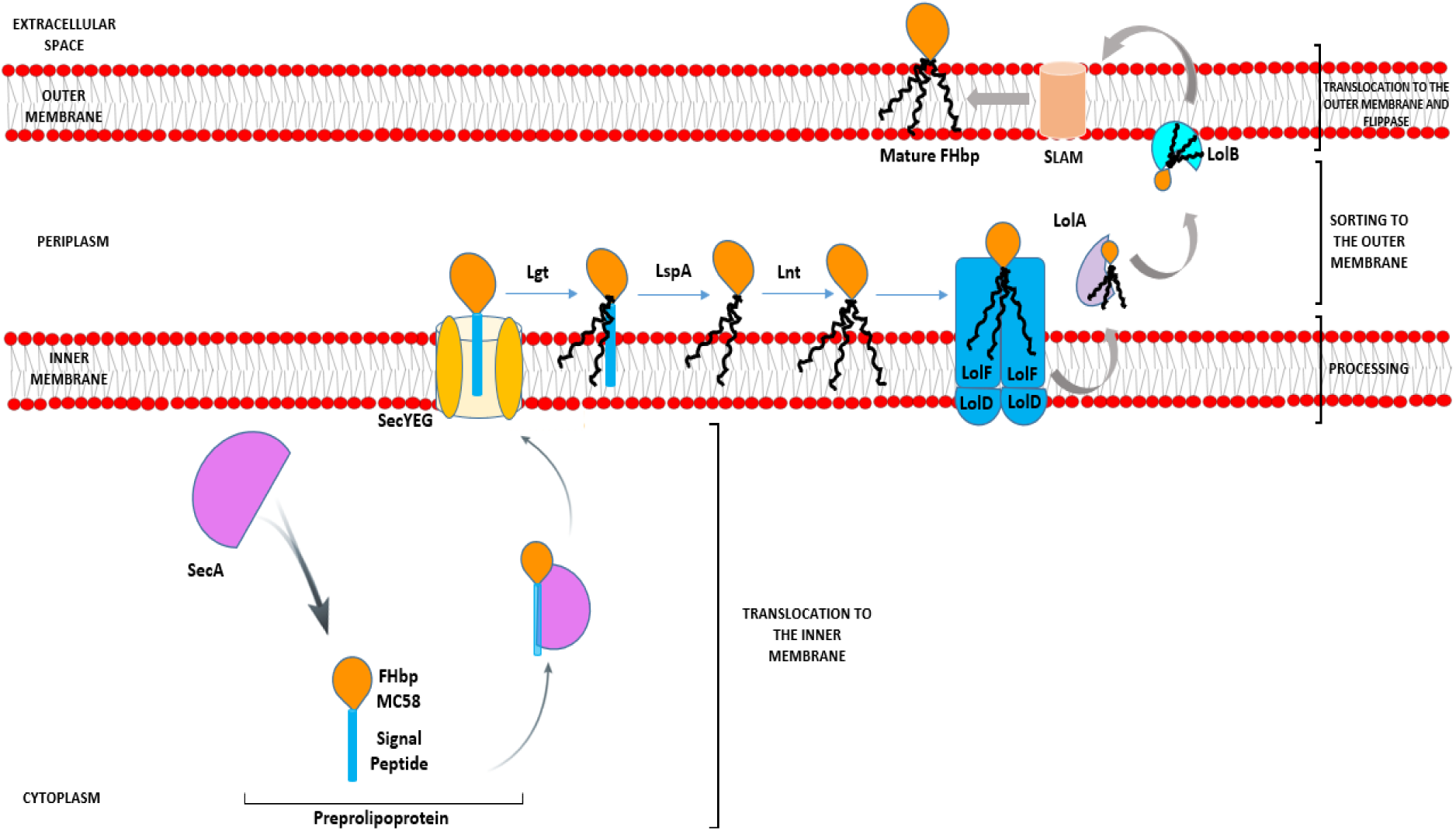
Model pathway for translocation, processing, sorting and export of FHbp in *N*. *meningitidis*. Preprolipoproteins are synthesised in the cytoplasm and as they emerge from ribosomes bind chaperones such as SecA, which translocate them across the IM via SecYEG. Lgt disacylates the preprolipoprotein at the conserved C residue of the lipobox of the SP, next LspA cleaves the SP exposing the C residue for further acylation by Lnt [9]. The Lol apparatus (LolFD) in *N. meningitidis*, [6] sorts the triacylated lipoprotein to the OM by delivering this to chaperone LolA, which releases the mature lipoprotein to the OM lipoprotein acceptor, LolB. SLAM then localises FHbp to the cell surface [18].

Through an accelerated approval process, both Trumemba (Pfizer) and Bexsero (GSK) were licensed by the FDA in 2015 for immunisation to prevent invasive disease by meningococcal group B in the United States in individuals 10 to 25 years of age. Trumenba comprises 2 recombinant FHbps, one from subfamily A, the other from subfamily B, both containing the lipid moiety found in the native protein [4, 19]. A recombinant non-lipidated form of FHbp from subfamily B is also one of the antigens of the Bexsero vaccine (GSK) [20] licensed for infants from 2 months of age in Europe in 2013 and now in Canada, Australia and several counties in South America [21].

A concern however for FHbp-based vaccines is the variation in expression level of FHbp between strains by over 15-fold and a threshold level of expression of at least 757 molecules of FHbp per cell is required for killing by FHbp-specific antibodies, meaning that isolates with low FHbp surface decoration fail to be effectively targeted [5, 22, 23]. The regulation of FHbp expression is influenced by external factors; at the transcriptional level by both iron and oxygen availability and at the translational level, by temperature [24–26]. Our work has focused on bacterial-cell-intrinsic molecular factors, which govern the processing and localisation of FHbp to the surface [9] (Fig 1) which is key for target recognition following immunisation with FHbp-based vaccines.

One of the studies correlating surface exposure of FHbp and susceptibility to killing by anti-FHbp antibodies was conducted by Newcombe and co-workers [27]. While strain L91543 displayed very little FHbp at the surface and demonstrated lack of killing by anti-FHbp sera, the well characterised MC58 strain demonstrated strong surface expression of FHbp and efficient killing by anti-FHbp sera [27]. We further observed a difference in size in FHbp between these 2 strains, despite their high amino acid (AA) sequence identity across the same length open reading frame (Fig 2a) [28]. The FHbp of strain MC58 is of the expected size; 27 kDa [29] and is known to be tri-palmitoylated at the C residue [30] whereas the FHbp of L91543 is about 3 kDa larger (Fig 2a) [28]. We identified 2 non-synonymous single nucleotide polymorphisms (SNP) in the h-region of the FHbp SP of L91543 that we hypothesised affects translocation and processing and ultimately localisation to the cell surface [28]. In this study, by restoring these SNPs we provide strong evidence to support our hypothesis.

**Figure 2.**
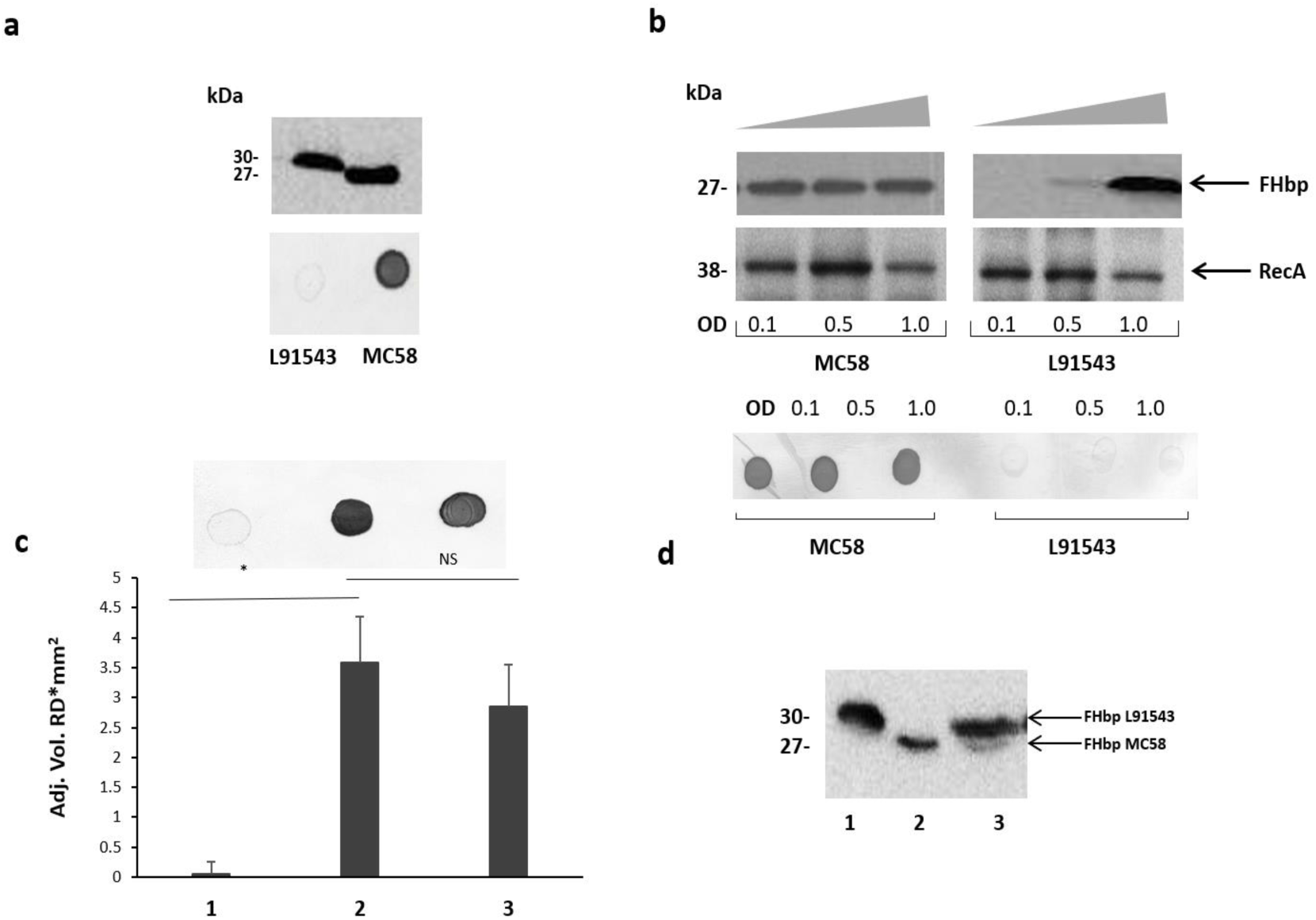
Cleavage and surface localisation of FHbp in L91543 upon transformation with MC58 *fHbp*. WCL (a, b and d) and whole cell suspensions from fresh plate cultures (a, b and c), or from broth cultures (b), analysed by Western immunoblot and immuno-dot blot respectively with JAR4. **a.** Western immunoblot (upper panel) and immuno-dot blot (lower panel) of strains L91543 and MC58. **b.** Different growth phases of broth cultures of MC58 and L91543 analysed by Western immunoblot (upper panel), including anti-recA antibody to verify loading control, and immuno-dot blot (lower panel), representative of 3 independent experiments. **c.** Immuno-dot blot and **d.** Western immunoblot: Lanes; 1, L91543; 2, MC58; 3, L91543*fHbp*MC58. Data are representative of 5 independent experiments. For **c**, the reflective density was measured by a GS-800™ calibrated densitometer. The one way ANOVA followed by Dunnett’s test was employed for statistical analysis in Graphpad (V. 6). All columns represent mean ± SEM, *p<0.05 vs strain MC58.

Surprisingly we found that the majority of invasive isolates circulating in the UK carry SP SNPs and exclusively express the preprolipoprotein which is partially retained in the cytoplasm due to reduced binding to SecA. Of the precursor protein that is translocated to the IM, some is retained in the periplasm and some exported to the surface. Our data suggest translocation to the OM is facilitated by Lnt and surface display is conducted by Slam following escape from processing by Lgt and LspA. Our findings thus suggest a novel role for Lnt as a chaperone facilitating periplasmic transport and for Slam in localising non-lipidated FHbp to the cell surface. We show that the reduced surface display of precursor FHbp, due to cytoplasmic retention, can be overcome by upregulating transcription. New algorithms can now be deployed to predict whether other preprolipoproteins are processed to become mature lipoproteins or remain as precursors. We investigated the biological implications of displaying precursor FHbp. Our findings bear relevance for vaccine efficacy data and we provide important insights for vaccinologists developing lipoprotein-based vaccines.

## Results

### SP SNPs in FHbp prevent cleavage but not surface localisation in strain L91543

*N. meningitidis* strain MC58 was originally isolated from a patient in the UK with meningococcal meningitis [31] and is a fully-sequenced, commonly used laboratory reference strain [32]. Strain L91543 was isolated from the cerebrospinal fluid of a 14-year old with meningococcal meningitis in the UK [33]. The FHbp-specific monoclonal antibody, JAR4, used throughout this study was sourced from the National Institute for Biological Standards and Controls (NIBSC). JAR4 binds to an N-terminal epitope including the AA residues DHK at positions 25 to 27 [34] (Table S1) and was generated following immunization of mice with MC58 FHbp [35]. The FHbps of both MC58 and L91543 are classified as subfamily B [4, 34] (variant 1) [6] and, as we previously reported, they share nucleotide and AA identities of 95 and 93% respectively [28].

**Table 1.**
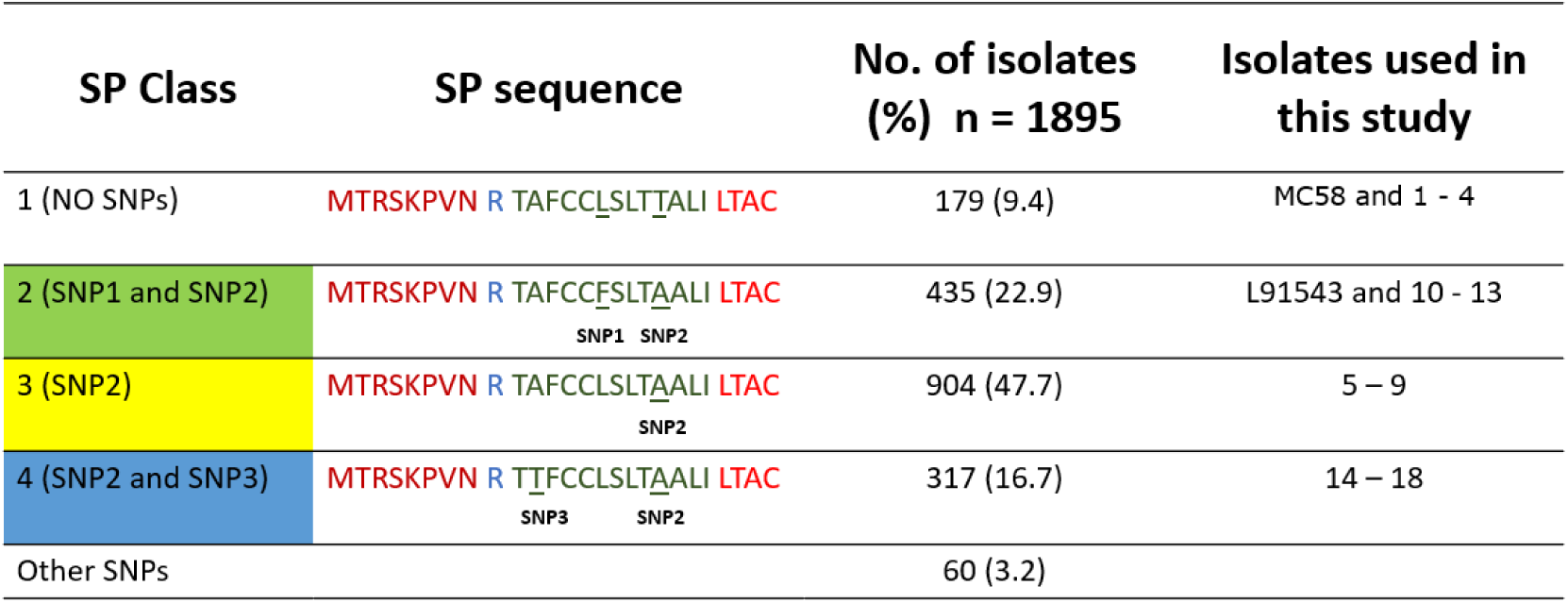
Frequency of different FHbp SP classes in capsular group B isolates in the MRF Meningococcus Genome Library database. The n-region is shown in brown, h-region in green and lipobox in red, corresponding to the display generated by the DOLOP lipoprotein prediction programme (https://www.mrc-lmb.cam.ac.uk/genomes/dolop/analysis.shtml) [11].

We investigated differences in the surface localisation of FHbp between these strains at different growth phases in a whole cell immuno-dot blot assay. A consistent strong level of surface expression of FHbp was shown for MC58 during growth in contrast to L91543, which demonstrated a uniform low level of expression (Fig 2a, 2b). Western immunoblot analysis of whole cell lysates showed a consistent high level of total FHbp expression for MC58, whereas the presence of FHbp was minimal in mid-log phase and strong only at stationary phase for L91543 cultures (Fig 2a, 2b). This demonstration of strong total FHbp level yet poor surface expression at stationary phase implies that FHbp is retained within the cells of L91543 and fails to either sort to the OM and/or become surface-exposed.

We previously reported that the FHbp of L91543 appears to be 3 kDa larger than that of MC58 upon fractionation of whole cell lysates as shown in Fig 2a, this additional mass corresponding to the size of the 26 AA SP [28]. To investigate any defects in LspA, which should cleave this SP, or aberrations in other proteins involved in the sequential pathway for FHbp processing and export [9] (Fig 1), the genome of L91543 was fully sequenced (GenBank accession number CPO16684) and BLAST analysis performed using relevant query sequences from MC58. LspA is 100% identical between the 2 strains and all other proteins share >97% identity with no evidence of frameshifts and premature stop codons, whereas FHbp exhibits only 93% AA identity between these 2 strains (Table S2). Notably the SP region of FHbp exhibits even greater divergence due to the presence of 2 SNPs in L91543; leucine (L) substituted by phenylalanine (F) at position 15 (SNP1) and threonine (T) substituted by alanine (A) at position 19 (SNP2) (Table 1).

**Table 2.** MenB invasive isolates used in this study. The SP class is indicated for each isolate.

| Class | Isolate Number | Isolate, full name |
| --- | --- | --- |
| 1 | - | MC58 |
|  | 1 | H44/76 |
|  | 2 | M10_240684 |
|  | 3 | M10_240701 |
|  | 4 | M02_241729 |
| 2 | 5 | M10_240579 |
|  | 6 | M13_240525 |
|  | 7 | M04_241215 |
|  | 8 | M11_241066 |
|  | 9 | M13_240614 |
| 3 | - | L91543 |
|  | 10 | M10_240750 |
|  | 11 | M13_240675 |
|  | 12 | M11_240236 |
|  | 13 | M12_240006 |
| 4 | 14 | M11_241033 |
|  | 15 | M14_240367 |
|  | 16 | M02_240210 |
|  | 17 | M11_240077 |
|  | 18 | M13_240486 |

To test our hypothesis and determine whether differences in FHbp sequence are solely responsible for the failure to cleave and export FHbp in strain L91543, the *fHbp* gene including its upstream promoter and ribosomal binding site from MC58 was inserted into the chromosome of strain L91543. This region was first cloned into Neisseria complementation vector, pGCC4, which allows integration of DNA in the intergenic region between *aspC* and *lctP* without causing polar affects [9, 36]. Plasmid pGCC4*fHbp*MC58 was transformed into strain L91543 and JAR4 was used to assess FHbp expression. Immuno-dot blot of whole cells showed that the recombinant strain, L91543*fHbp*MC58, acquired the ability to express FHbp at the cell surface (Fig 2c). Western immunoblot analysis of L91543*fHbp*MC58 revealed 2 bands: a lower band with a mobility equivalent to the cleaved FHbp of MC58 and a higher molecular weight band corresponding to the expected mobility for the native, uncleaved protein (Fig 2d).

To investigate the specific and combined contribution of the 2 individual SNPs in L91543 SP on cleavage and export, first *fHbp* was deleted in L91543 (L91543Δ*fHbp*) then complemented with wild type (L91543Δ*fHbp*+L*fHbp*) or versions with site-directed alteration in the SP (L91543Δ*fHbp*+L*fHbp*SNP1, L91543Δ*fHbp*+L*fHbp*SNP2 and L91543Δ*fHbp*+L*fHbp*SNP1+2). Western immunoblot analysis confirmed complete loss of FHbp expression in L91543Δ*fHbp* (Fig S1). L91543Δ*fHbp*+L*fHbp*SNP1 and L91543Δ*fHbp*+L*fHbp*SNP2 showed moderate levels of surface expression and L91543Δ*fHbp*+L*fHbp*SNP1+2 showed a level of surface display comparable to MC58 (Fig 3). Interestingly Western immunoblot analysis showed higher mobility, indicative of SP cleavage, for only strain L91543Δ*fHbp*+L*fHbp*SNP1+2, which had both SNPs corrected but not for either recombinant strain with individual SNPs corrected (Fig 3). Our results suggest that both AA substitutions impact on the FHbp translocation-processing pathway at some point up to, or at, the SP cleavage step and surprisingly that repair of either SNP individually can permit partial localisation of FHbp to the cell surface without the need for prior SP cleavage.

**Figure 3.**
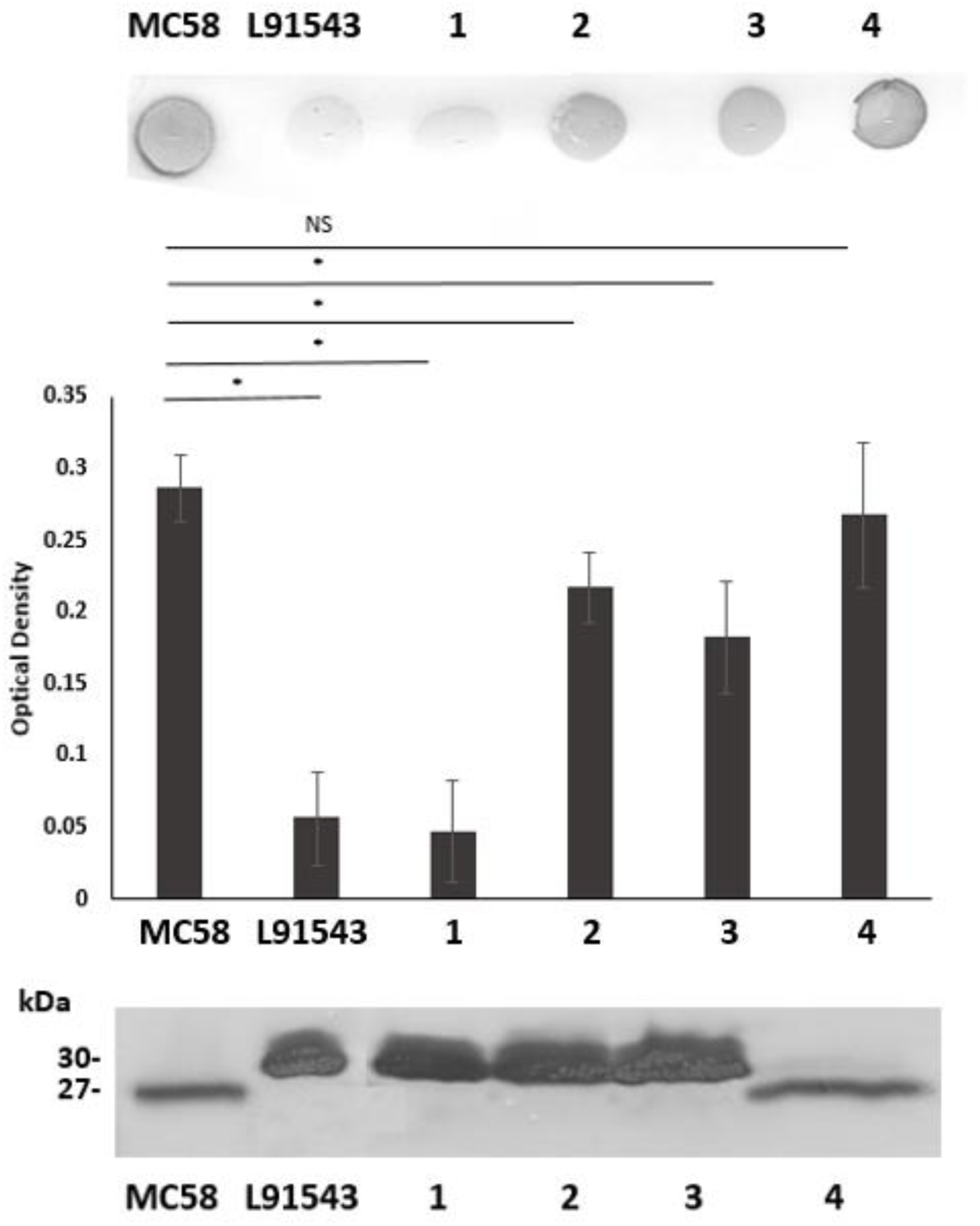
SP SNPs in FHbp prevent cleavage but not surface localisation. Whole cell suspensions and WCL from fresh plate cultures analysed by immuno-dot blot (upper panel) and Western immunoblot (lower panel) respectively with JAR4. Lanes; 1, L91543Δ*fHbp+*L*fHbp*; *2,* L91543Δ*fHbp*+L*fHbp*SNP1; 3, L91543Δ*fHbp*+L*fHbp*SNP2; 4, L91543Δ*fHbp*+L*fHbp*SNP1+2, representative of 3 independent experiments. The density of spots was measured by ImageJ and the one way ANOVA followed by Dunnett’s test was employed for statistical analysis in Graphpad (V. 6). All columns represent mean ± SEM, *p<0.05 vs strain MC58.

### FHbp with SP SNPs shows reduced binding to SecA

Most proteins destined for the OM are first targeted to the IM translocase, Sec-YEG, by the chaperone ATPase motor, SecA. SecA recognizes SPs by having a high affinity for their hydrophobic binding grooves and by electrostatically trapping their positively charged n-region through acidic residues [37, 38]. We investigated the binding of L91543 FHbp (carrying SNP1 and SNP2) to meningococcal SecA and compared this to MC58 FHbp, which is known to undergo cleavage and processing [6, 9]. The bacterial 2-hybrid approach was taken which relies on induced expression of the 2 heterologous proteins using *E. coli* as the host organism. The cytoplasmic readout for binding affinity in this heterologous expression system is expected to be similar to that in the meningococcus, prior to translocation to the IM. Our results showed a significant 1.9-fold lower binding of SecA to L91543 FHbp as compared with MC58 FHbp (Fig 4).

**Figure 4.**
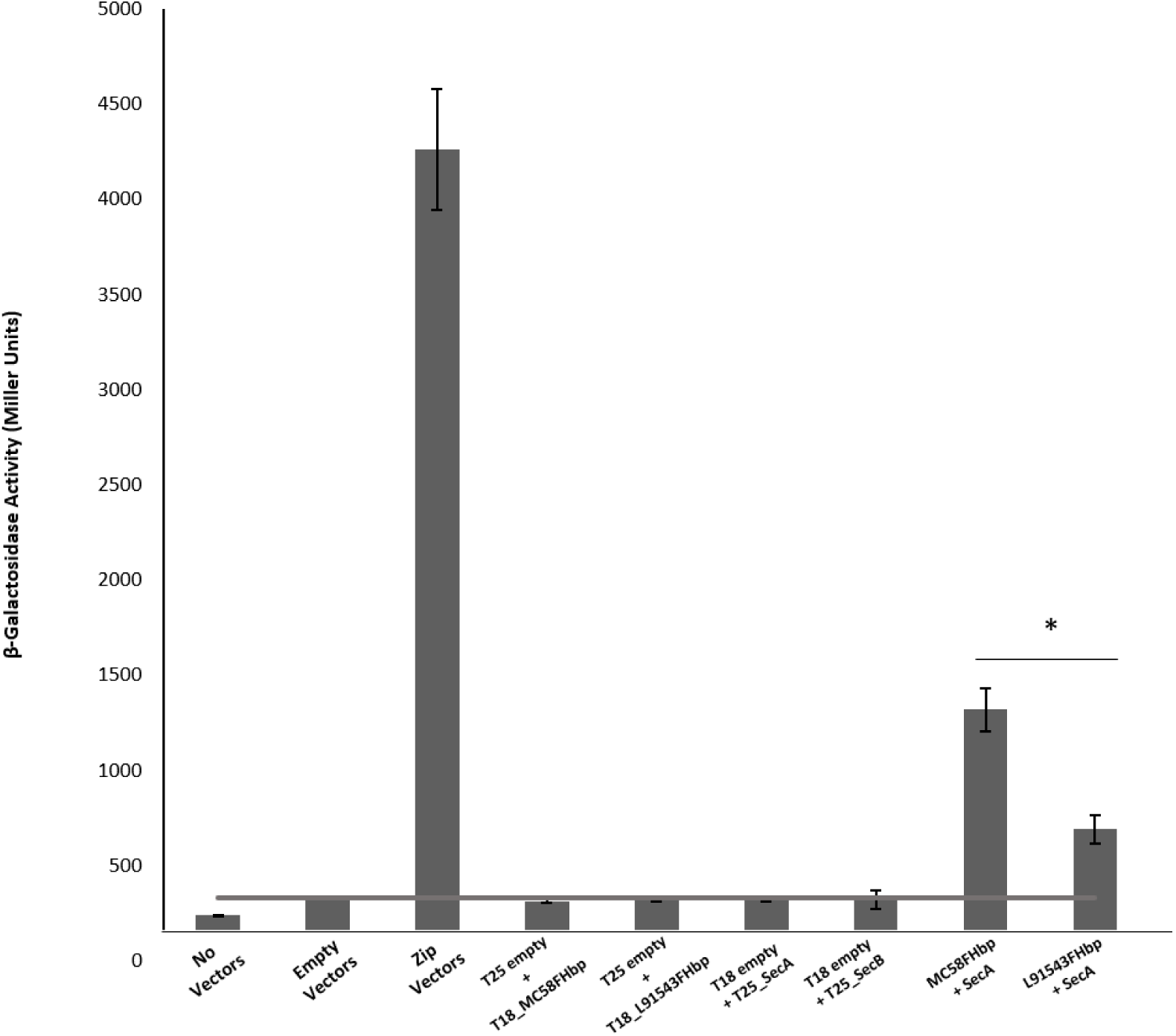
FHbp with SP SNPs shows reduced binding to SecA. Bacterial 2 hybrid experiments to compare FHbp binding to SecA in MC58 and L91543. The *fHbp* gene of strains MC58 and L91543 and *secA* gene of MC58 were cloned into the two-hybrid pUT18 (T18-prey) and pKT25 (T25-bait) vectors respectively and the appropriate prey and bait pair of vectors co-transformed in BTH101. In parallel, zip prey and zip bait vectors were co-transformed in BTH101 as a positive control for protein-protein interaction. Likewise, different prey-bait combinations but with one vector or both vectors empty were used as negative controls. The interactions were quantified by measuring the ß-galactosidase level expressed in Miller units (values taken from 4 independent clones). Unpaired T Tests with Welsch’s correction were conducted to compare the interactions. All columns represent mean ± SEM, *p<0.05 vs MC58 FHbp.

### Clinical isolates with FHbp SP SNPs show SP retention yet surface localization

We next investigated the prevalence of FHbp containing SP SNPs that differ from the MC58 sequence in the genome sequences of 1,895 invasive group B UK isolates in the Meningitis Research Foundation (MRF) Meningococcus Genome Library database collected between 2009 and 2017. Unexpectedly we found that only 9% of isolates have a FHbp SP identical to that of MC58 (referred to as class 1) with the remaining isolates carrying either (i) both SNP1 and SNP2, like L91543 (class 2, 23%); (ii) SNP2 alone (class 3, 48%); (iii) SNP2 and a third SNP (SNP3) (class 4, 17%), or (iv) other SNPs (3%) (Table 1). All SNPs are located exclusively in the h-region 13 AA in length, and common to class 2, 3 and 4 is SNP2 (i.e. T substituted by A at position 19).

To test if our findings for recombinant strains of L91543 apply to these MRF isolates with non-class 1 FHbp, a selection of isolates from each class were randomly selected (Table 2). Strain H44/76 was chosen as an additional positive control strain since its FHbp including SP is identical to that of MC58. FHbps of all 20 isolates belong to subfamily B/variant 1 (the most prevalent group) and bear the JAR4 epitope [34] (Table 2, Table S1).

FHbp from all isolates with class 1 SP demonstrated a similar size by SDS-PAGE to that of MC58 whilst all non-class 1 FHbp proteins had the lower mobility observed for L91543 (Fig 5a). To determine whether the change in mobility affected surface localisation, whole cell immuno-dot blots were performed with JAR4 except for isolate 4 which JAR4 failed to recognise. For this isolate, polyclonal anti-FHbp sera was used. The median level of surface expression was highest for class 1 isolates and decreased progressively from class 3 isolates (SNP2 only), to class 4 isolates (SNP2 and SNP3) to class 2 isolates (SNP1 and SNP2) (Fig 5b). Overall, non-class 1 isolates demonstrated 2.2-fold lower surface expression of FHbp than class 1 isolates (excluding isolate 4) with p 0.0037 (Fig 5b). These results indicate a failure to cleave the SP decreases the efficiency but does not abrogate localistion of FHbp to the surface. This data support our findings for recombinant strains of L91543 (Fig 3). In contrast to the general pattern, 3 isolates (L91543, 12 and 13) demonstrated the presence of FHbp by Western immunoblot but no localization to the cell surface by immuno-dot blot (Fig 5b), the cause of which is investigated later in this manuscript.

**Figure 5.**
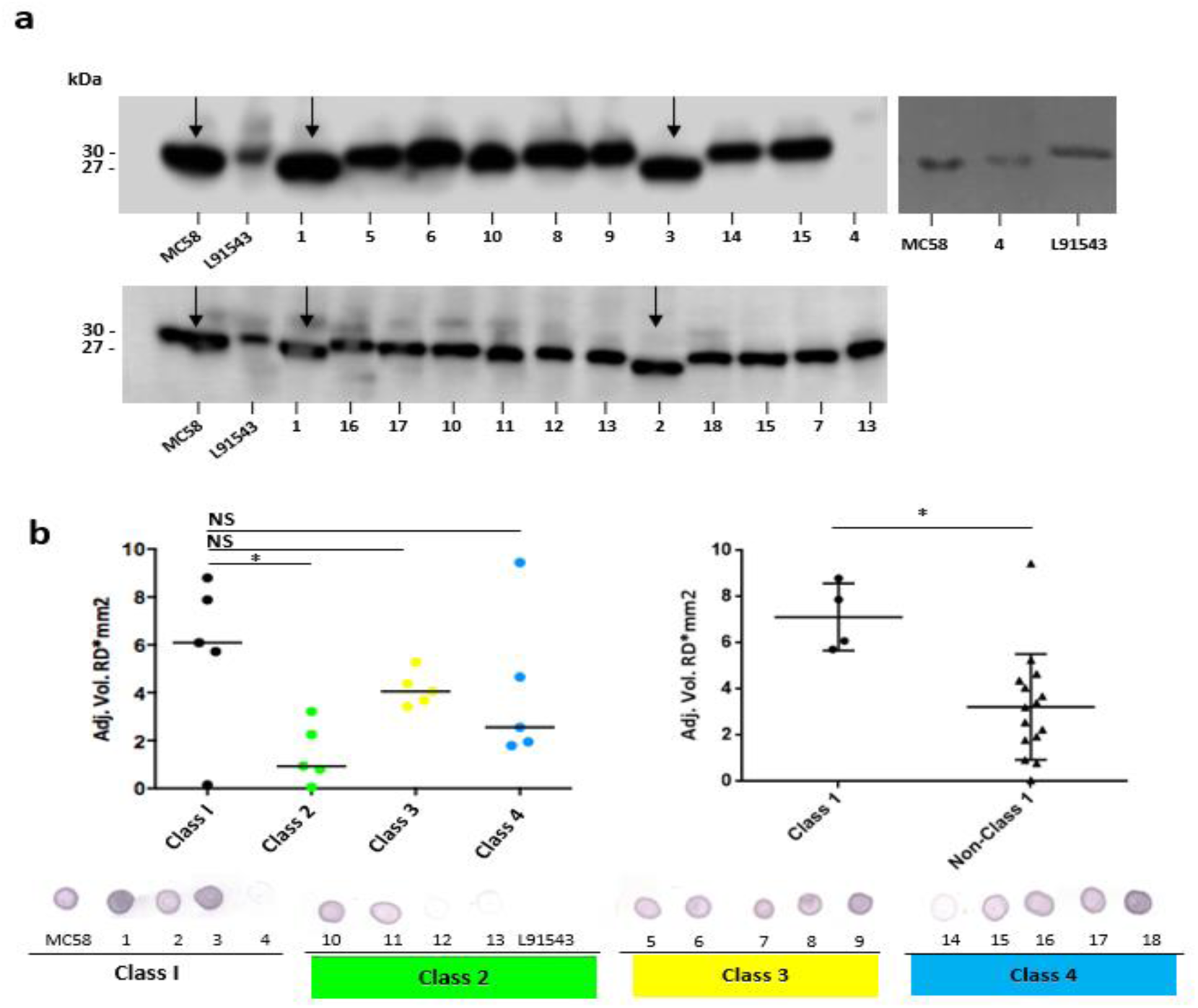
Comparison of FHbp size and surface localisation in clinical isolates. For all non-class 1 isolates, the SP is uncleaved yet for most of these isolates, FHbp is surface localised. WCL or whole cell suspensions were prepared from fresh plate cultures and JAR4 used except where indicated. **a.** Western immunoblot. For isolate 4, polyclonal anti-FHbp antibody was also used; **b**, Whole cell immuno-dot blot. The density of spots was measured by ImageJ and the one way ANOVA followed by Dunnett’s test employed for statistical analysis in Graphpad (V. 6). All columns represent mean ± SEM, *p<0.05 vs class 1. Pooled data of class 1 and non-class 1 were analysed using un-paired T Tests with Welsch’s correction. Both columns represent mean ± SEM, *p<0.05 vs class 1.

Interestingly all strains shown to express uncleaved FHbp in this study carry SNP2: substitution of the polar T residue with a hydrophobic A residue at position 19. The importance of polar residues in the h-region of the SP of bacterial lipoproteins first came to light in the late 1970s. In all but one of the 26 bacterial SPs known at that time, polar residues; serine (S) and/or T were found in the h-region and within close proximity to the signal peptidase cleavage site [39]. A T16A mutant within the SP of *E. coli* Lpp lipoprotein affected the rate of processing with accumulation of unmodified precursor in the membrane fraction [38]. It was postulated that this T residue (as well as a serine at position 15) either directly affected recognition by Lgt or LspA or affected IM translocation hindering accessibility of the SP by these enzymes. We questioned whether the meningococcal Lgt and LspA enzymes fail to recognise FHbp with class 2 (SNP1 and SNP2), class 3 (SNP2) and class 4 (SNP2 and SNP3) SPs (Table 1) and how these preprolipoproteins localise to the surface.

### FHbp with SP SNPs localise to the surface via Lnt and Slam with escape from processing by Lgt and LspA

To test if strains with SP SNPs in FHbp are affected in lipid modification, one isolate representative of each SP class was tested for the presence of the lipid moiety. These included MC58 (class 1), L91543 (class 2), isolate 6 (class 3) and isolate 18 (class 4) (Table 1 and 2). The strains were grown in medium containing palmitic acid alkyne, FHbp was immuno-precipitated then conjugated (clicked) with biotin azide, which attaches exclusively to proteins carrying this alkyne. Western immunoblot analyses were performed separately with streptavidin and with JAR4. A band of the expected mobility for cleaved FHbp was detected by streptavidin for MC58 clicked with biotin azide and not for the corresponding non-clicked control (Fig 6a). However, no bands were detected for L91543, isolate 6 or 18 whether clicked or non-clicked, suggesting that the FHbp molecules of these 3 isolates are not lipidated.

**Figure 6.**
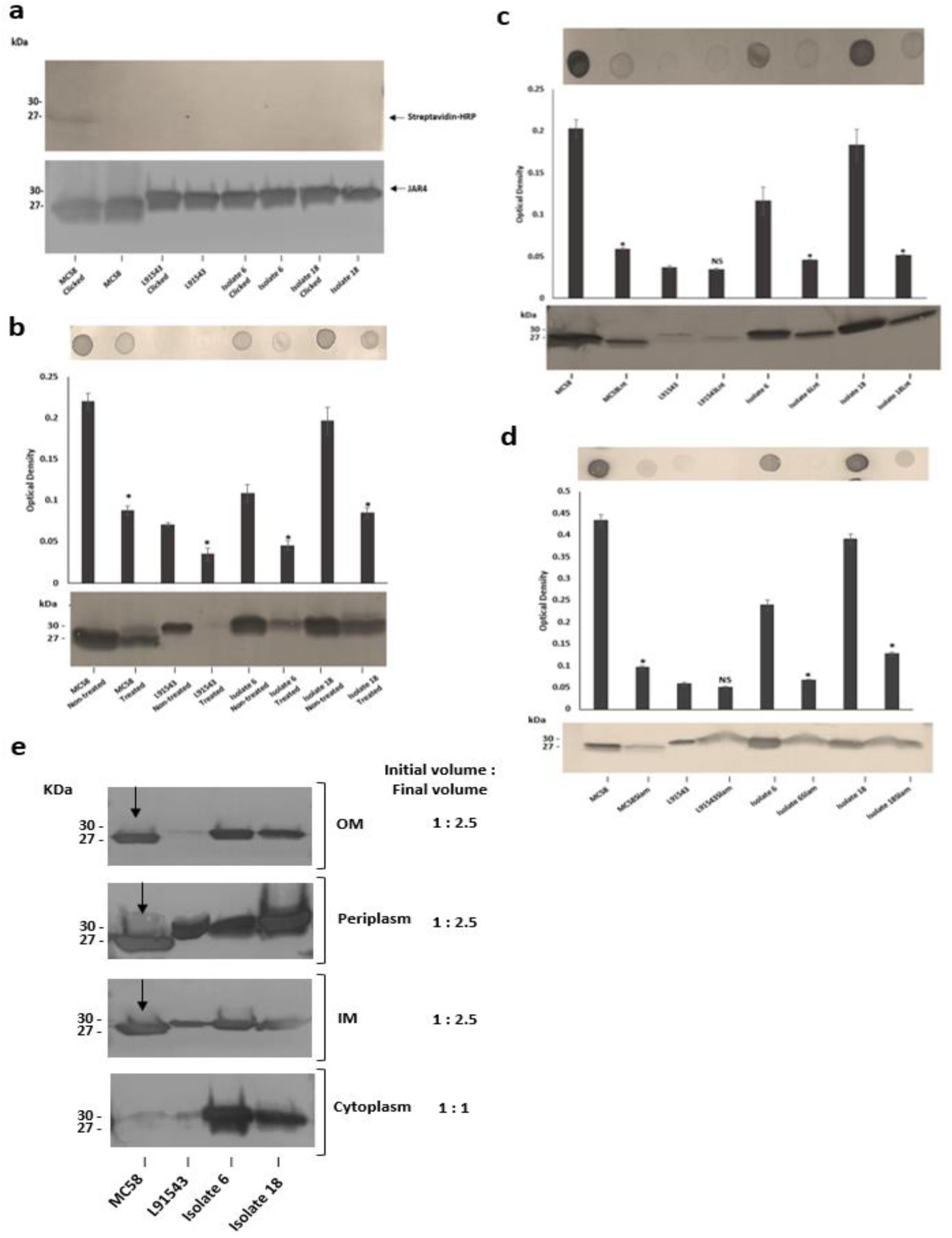
FHbp with SP SNPs localise to the surface via Lnt and Slam with escape from processing by Lgt and LspA. **a.** Western immunoblot after growing strains in palmitic acid alkyne and clicking immuno-precipitated FHbp with biotin-azide; JAR4 (upper panel) and Streptavidin-HRP (lower panel). **b to d.** Immuno-dot blot of broth cultures (*A*_600_ 0.5) and Western immunoblot of WCL with JAR4, (b, after globomycin treatment). Unpaired T Tests with Welsch’s correction were conducted to compare the isogenic strains. All columns represent mean ± SEM, *p<0.05 vs wild type isogenic strain. **e**. Western immunoblot of different cellular fractions with JAR4. (Lane 1, MC58; 2, L91543; 3, isolate 6; 4, isolate 18.)

To test if substrate recognition by LspA for non-class 1 FHbp (non-lipidated) is affected, the isolates were incubated with and without globomycin which specifically inhibits LspA [18] and lysates were analysed by Western immunoblotting. For MC58, the detection of both the 30 kDa and 27 kDa bands suggests partial inhibition of FHbp processing by LspA (Fig 6b). For L91543 and isolates 6 and 18, there was no change in their FHbp mobility; each displayed the 30 kDa form, demonstrating that inhibition of LspA had no affect on FHbp (Fig 6b). The reduced level of FHbp for all 4 globomycin-treated strains was not unexpected given the global impact LspA has on the entire lipoproteome including LolB, directly involved in OM translocation of lipoproteins [17].

Following failure of Lgt and LspA to lipidate and process FHbp in the non-class 1 strains, we investigated whether the canonical pathway is terminated from this point onwards or resumed by Lnt fur subsequent surface localisation. The *lnt* gene of the 4 strains was disrupted. Other than L91543 parent and mutant strain, which both showed the typical low level of FHbp by Western immunoblotting and immune-dot blotting, the Lnt mutant of the other 3 strains showed a significant reduction in surface localisation as well as a diminished level of total FHbp (Fig 6c). This supports our previous study where we showed that MC58Lnt employs mechanisms to reduce FHbp levels thereby preventing the toxic accumulation of the diacylated lipoprotein in the periplasm by down-regulating FHbp expression and likely employing periplasmic proteases as well as shunting some of this FHbp to the surface [9]. The observation of a similar reduction in FHbp level for isolates 6 and 18 suggests Lnt plays a role in FHbp periplasmic transport regardless of whether FHbp is processed by LspA and Lgt or not.

Given that the canonical pathway appears to resume from Lnt, we investigated whether Lol was employed to translocate FHbp of these strains to the OM however it was not possible to mutate any of the Lol-encoding genes. We next investigated if Slam plays a role in the surface localisation of FHbp in these strains. Again with the exception of the poor FHbp expressor, L91543, the disruption of Slam resulted in significantly reduced surface localisation of FHbp as well as some reduction in total FHbp (Fig 6d). This suggests that Slam plays a role in surface localisation of FHbp regardless of whether FHbp is processed or not. This contradicts what was proposed by Hooda et al., (2016) that Slams are responsible for transport of surface lipoproteins that possess a lipid anchor that needs to be flipped from the inner leaflet to the outer leaflet of the OM [18]. Rather, in this study we show that surface localisation by Slam is not restricted to lipoproteins.

To confirm that these alterations in FHbp phenotypes were not an artefact of introducing mutations per se into the chromosome, 2 different genes encoding pili (*pilE* and *pilS*) were deleted in all 4 strains and no alteration in FHbp mobility, quantity or surface localisation was detected (data not shown).

Given our findings that FHbp with SP SNPs binds less well to the IM translocase, SecA, but appears to be translocated to the OM and surface localised by Slam, we would expect this to be reflected in the sub-cellular distribution of FHbp for these 4 strains. In MC58 a processed FHbp protein of size 27 kDa was found in the IM, periplasm and OM but was barely detectable in the cytoplasm, providing evidence for highly efficient translocation from cytoplasm to the IM but less efficient translocation to the OM (Fig 6e). In contrast, for isolates 6 and 18, unprocessed precursor was detected in all 4 cellular compartments; cytoplasm, IM, periplasm and OM, and at high levels (Fig 6e). These findings demonstrate that the efficiency of translocation from cytoplasm to IM is reduced in these isolates (supporting our BACTH data) but surprisingly their OM translocation efficiency is comparable with that of MC58. For strain L91543, unprocessed FHbp was noticeably less abundant and all of this localised to the IM and periplasm suggesting complete translocation across the IM (Fig 6e). Conversely, for isolates 6 and 18, the greater overall abundance of preprolipoprotein appears to have exceeded the capacity of SecA to fully translocate it. From this, we propose that when the FHbp preprolipoprotein is abundant, SecA which is already hampered in its binding affinity for non-class 1 SPs, translocates only a portion of this precursor to the IM.

### Transcript levels of *fHbp* influences surface localisation of FHbp

To explore whether differences in the level of transcription of *fHbp* in the isolates accounted for the variations in level of FHbp preprolipoprotein at the cell surface, RT-PCR with invariant *fHbp*-specific primers was performed from *A*600 1.0 broth cultures. All isolates demonstrated a similar level of *fHbp* transcript except for the 3 isolates that displayed FHbp poorly at the cell surface (Fig 7). These 3 isolates exhibited 4 to 6-fold reduction in *fHbp* transcripts (Fig 7).

**Figure 7.**
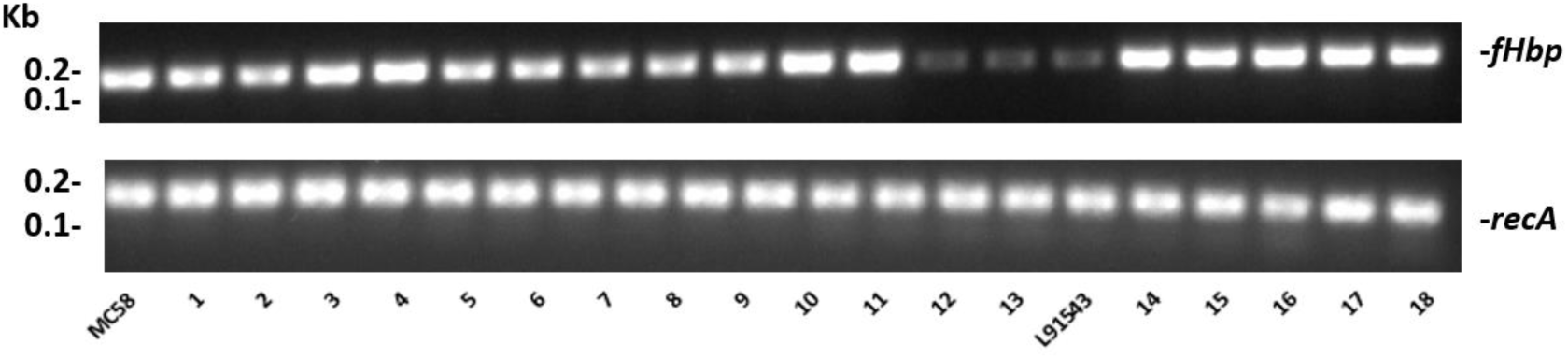
Comparison of *fHbp* transcript level between isolates. RT-PCR of *fHbp* and recA (standard house-keeping gene). One representative experiment of 3 is shown.

To test if increasing the rate of transcription promotes the localisation of precursor FHbp to the surface, we utilised strain L91543Δ*fHbp*+L*fHbp* wherein the native class 2 *fHbp* is under the control of a *lacZ* promoter. The amount of surface FHbp preprolipoprotein was compared by immuno-dot blot between isopropyl-β-D-thiogalactopyranoside (IPTG)-induced and non-induced L91543Δ*fHbp*+L*fHbp*, relative to the L91543 parent strain and to MC58. The induced recombinant strain localised unprocessed FHbp to the surface with similar levels as observed for MC58 (Fig 8a).

**Figure 8.**
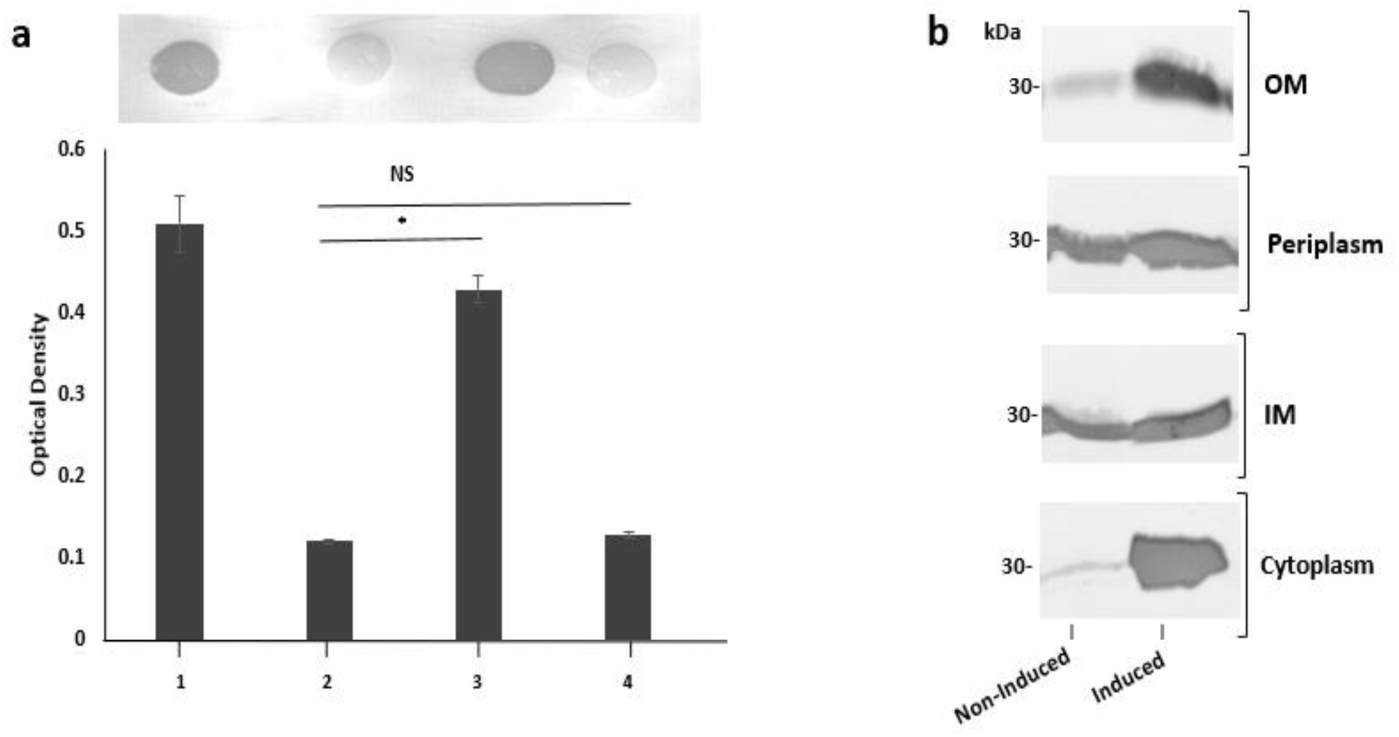
The affect of increasing transcription of *fHbp* on FHbp subcellular localization and surface localization in L91543. **a.** Immuno-dot blot of whole cells with JAR4. Lanes; 1, MC58, 2, L91543, 3, L91543Δ*fHbp*+L*fHbp* induced with IPTG, 4, L91543Δ*fHbp*+L*fHbp* non-induced. One representative experiment of 5 is shown. The optical density was measured by ImageJ. The ANOVA followed by Dunnett’s test was employed for statistical analysis in Graphpad (V. 6). All columns represent mean ± SEM, *p<0.05 vs strain L91543; **b.** Western immunoblot with JAR4 of different subcellular compartments of strain L91543Δ*fHbp*+L*fHbp* non-induced versus induced.

Next we investigated the subcellular distribution of precursor FHbp in these recombinant strains. In the absence of induction, the preprolipoprotein localized to the periplasm and IM but very poorly to the OM (Fig 8b) similar to that for the L91543 parent strain (Fig 8b). Upon induction, not only did the quantity of preprolipoprotein dramatically increase, as expected, but as with isolates 6 and 18, some was retained in the cytoplasm while the remainder localized to the IM, periplasm and OM (Fig 8b). Together the data suggest that a certain threshold level of FHbp in the periplasm is tolerated, as shown for strains L91543 (Fig 6e) and non-induced recombinant strain, L91543Δ*fHbp*+L*fHbp* (Fig 8b). However, over-accumulation in the periplasm appears to force translocation to the OM as observed for induced L91543Δ*fHbp*+L*fHbp* and for isolates 6 and 18 (Fig 8b).

### Comparison of biological activities of unprocessed and processed surface FHbp

Crystal structure studies reveal that FHbp folds to form an N-terminal β barrel and a C-terminal β barrel and that the FHbp-fH complex is held together by extensive interactions between both β-barrels and domains 6 and 7 of fH [29]. We questioned whether the inclusion of the 26 AA SP that is predicted to form an α-helix at the N terminus of FHbp affects the binding of FHbp to fH. Equivalent cell numbers of each of the 20 isolates were incubated with fH. Anti-fH antibody binding was then measured by Fluorescence-Activated Cell Sorting (FACS) and no significant difference in binding of class 1 versus non-class 1 isolates was found.

Since a certain threshold level of FHbp on the surface of cells is required for successful killing by vaccine-induced antibodies [5, 40] we questioned whether isolates with non-class 1 FHbp SP that typically display less unprocessed FHbp at the surface may be more resistant to killing by antibodies. First FHbp surface expression was quantified by FACS analysis with JAR4. Since this monoclonal antibody binds a single epitope (Table S1), saturation of meningococcal cells with this antibody should result in the maximal binding of one antibody per FHbp molecule. Commercially available microspheres were used to quantify the number of antibodies bound per cell (ABC) and enable us to infer from this the number of FHbp molecules displayed per cell. The SBA assay was performed using the FHbp-specific monoclonal antibody JAR5 along with JAR4. Cooperativity between these 2 antibodies is necessary to elicit bactericidal activity in the presence of human complement [41–43]. JAR5 is known to bind the FHbp epitope comprising AA residues 84-91 and 115-123 [41] (Table S1). Strains MC58 and L91543 were previously shown to contrast starkly in their FHbp exposure and susceptibility to killing by FHbp-immunised sera in SBA assays [27] and were used as positive and negative control strains respectively to benchmark all other isolates.

We first noted that isolates 4, 12 and 13 displayed very poor surface FHbp at a level comparable with that of L91543 (Fig 9a) supporting our previous immuno-dot blot data for these isolates (Fig 5b). These levels were well below 757 molecules per cell, the threshold level for killing by bactericidal anti-FHbp antibodies [23]. Unsurprisingly, extremely weak bactericidal activity was detected for these isolates in SBA assays with only up to 11% killing demonstrated (Fig 9b). Our findings are explained by the poor transcript levels observed for isolates L91543, 12 and 13 (Fig 7) and by failure of JAR4 to bind isolate 4 (Fig 5b). Pooled class 1 isolates were compared with pooled non-class 1 isolates after discounting the 4 outliers. For ABC, the average was 7869 vs 4547 molecules (p=0.0038) and for SBA the average was 73.7% vs 53.4% (p=0.0009) (Fig 9c). We anticipate seeing greater differences in SBA responses between these 2 groups using sera from Trumenba-vaccinated individuals since this vaccine targets cleaved, lipidated FHbp, but we were unable to obtain this sera. Such sera will contain diverse polyclonal antibodies that bind to multiple epitopes of FHbp.

**Figure 9.**
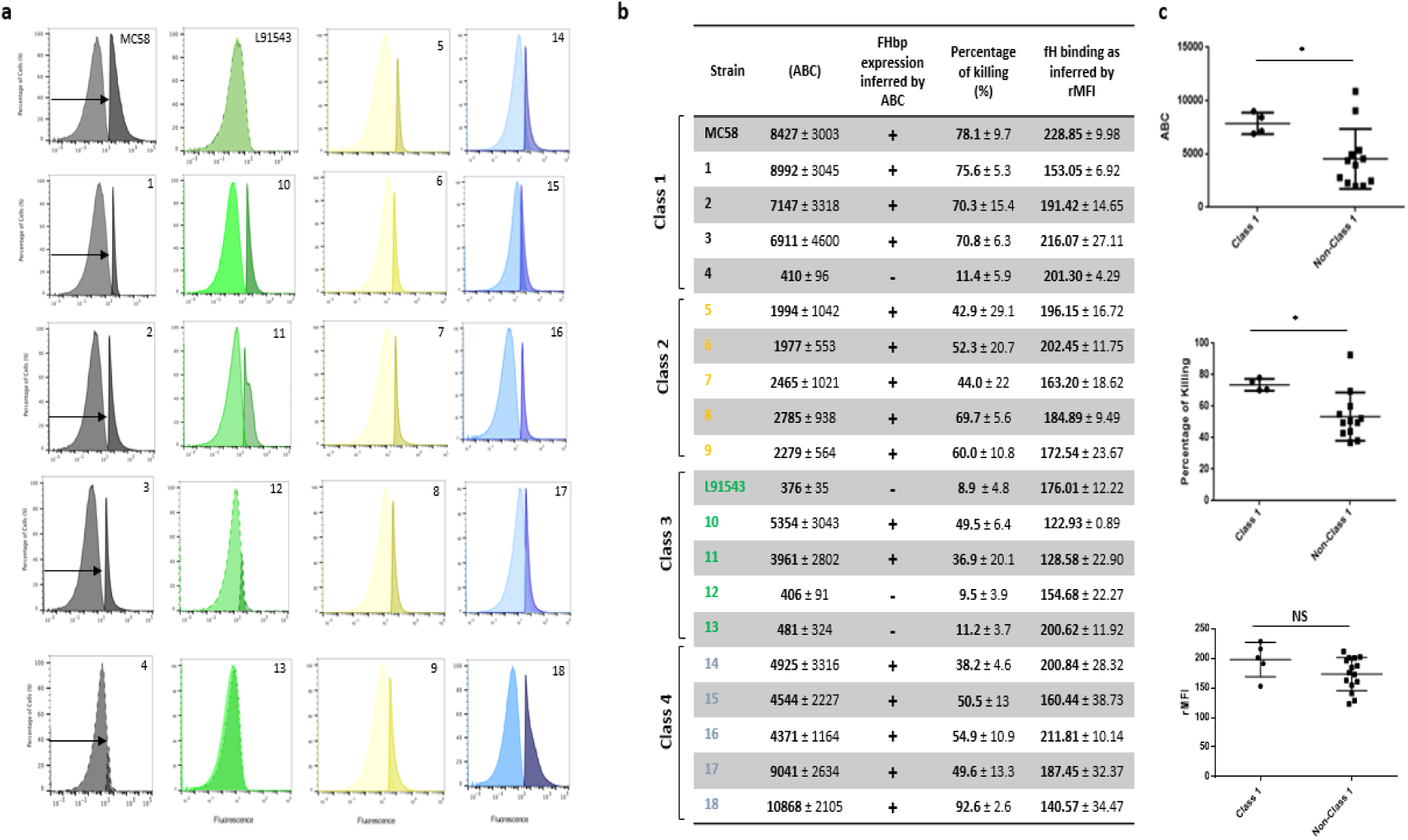
Comparison of biological activities of unprocessed and processed surface localised FHbp. **a.** FACS analysis with Mab JAR4. A representative flow cytometry plot for each isolate is shown. The read-outs from negative control samples, cells incubated with secondary antibody only, (left peak) were gated (as shown by the arrows). The read-outs from samples incubated with both primary and secondary antibody were overlaid; **b.** The number of JAR4 antibody molecules bound per cell (ABC) and corresponding prediction for successful killing in SBA assays is denoted by +. The actual percentage killing for each isolate with Mabs JAR4, JAR5 and human complement is shown. The mean ABC for each isolate was derived from 3 independent FACS experiments and the mean percentage killing derived from 4 independent SBA experiments, each with 2 technical replicates. The binding of fH to cells. Values show the Relative Mean of Fluorescence Intensity (rMFI) for total fluorescing cells after incubation with fH and anti-fH PE-conjugated antibody. The values were normalised against the negative control of cells alone incubated with antibody. **c.** Pooled data for class 1 and non-class 1 isolates, excluding the outliers. Values were analysed by unpaired T Tests with Welsch’s correction. Columns represent mean ± SEM, *p<0.05 vs class 1.

## Discussion

Since the discovery of FHbp as a lipoprotein [4] it has been assumed that all meningococci that express FHbp synthesise the mature lipoprotein. The loss of processing, including SP cleavage and lipidation, in strain L91543 was directly linked to mutations in the SP that interrupt Sec-mediated translocation. The correlation between SP mutations and failure to process FHbp was further confirmed in a panel of invasive group B isolates [44]. Remarkably the preprolipoprotein localised to the cell surface but with over 2-fold lower abundance and in the subset of isolates tested, this was attributed to some retention in the cytoplasm. For L91543 and likely the remaining minority of isolates with poor transcription strength of *fHbp*, the less abundant preprolipoprotein was all translocated across the IM but none was translocated to the OM for subsequent surface display. This could however be overcome by increasing the transcription strength of *fHbp*.

It is known that Sec-dependent proteins are exported in unfolded states to the Sec-YEG channel (Fig 1) as this channel is only large enough to accommodate unfolded proteins [45]. SecA acts promptly to bind nascent polypeptides as they emerge from the ribosome during translation and maintains the polypeptide in an unfolded state [46]. Disrupting this interaction causes a partial defect in SecA-mediated translocation [46] and reduced efficiency of transport to the IM due to the propensity for the preprolipoprotein to fold and become incompetent for translocation [47]. Based on new insights in SecA-mediated translocation [48], it is possible that the preprolipoprotein that is translocated is not orientated correctly in the IM such that Lgt fails to recognise this FHbp precursor as substrate.

We propose that the presence of a polar AA (in this case, T) four AAs away from the lipobox of FHbp, is the most critical for correct translocation by SecA as shown for MC58 and all other class 1 isolates tested. All non-class 1 isolates showed lack of processing and have the T19A mutation and for those with class 3 SP, T19A is the sole SP mutation. This mutation is also the sole mutation in 48% of MRF isolates and is represented in at least 88% of MRF isolates (Table 1). We postulate that in T19A FHbp variants, the reduced efficiency of binding to the SP by SecA results in the propensity for folding with consequent retention of some of this translocation-incompetent preprolipoprotein in the cytoplasm. This retention was evident for both isolates 6 and 18 tested that express an abundance of the preprolipoprotein and for L91543 after induction of strong expression of FHbp (Fig 6e, Fig 8b). The remaining preprolipoprotein was translocated across the IM and some of this was translocated to the OM and further surface-displayed. Upon accumulation of the unprocessed precursor FHbp in the IM beyond a threshold level, translocation to the OM is forced, probably to prevent clogging which would prohibit the translocation of other OM-destined proteins and be potentially fatal for the cell. We provide data suggesting this translocation is partly mediated by Lnt highlighting a secondary role for Lnt, possibly as a chaperone facilitating delivery to Lol. We further show that localisation to the surface involves Slam revealing that Slam is not lipoprotein-specific. Slam thus appears to be a surface “protein” assembly modulator in addition to a surface lipoprotein assembly modulator and has broader specificity for OM protein localisation than originally thought. This is plausible given Slam recognises the C terminal structure of its substrates [18] which may be shared between lipoproteins and OM proteins.

From analysis of the SP sequence of 1895 invasive group B isolates from the UK MRF collection, as few as 9% are predicted to express FHbp as a lipoprotein. Conversely, at least 88% are predicted to display the unprocessed, precursor at the surface, provided sufficient transcription strength of *fHbp*. Similar percentages are expected for other isolates circulating around the globe. Given that isolates displaying non-class 1 FHbp SP are far more prevalent than class 1 isolates, there must be a fitness benefit for the former. One explanation for evolutionary selection for the former could be that exportation of unprocessed FHbp is less metabolically costly than employing processing enzymes to modify FHbp prior to localisation of the mature lipoprotein to the surface. Alternatively, unprocessed FHbp at the surface may be more resistant to bactericidal killing in the host than the mature lipoprotein. The significantly reduced surface exposure of the preprolipoprotein, shown in this study, may be selected for to escape anti-FHbp antibody-mediated killing whilst retaining sufficient fH binding.

Our results reveal that the AA sequence requirement for lipoprotein SPs to direct translocation to the IM completely and correctly for processing to follow, are more constrained than previously thought. For FHbp translocation and maturation, the polar residue close to the lipobox is critical and this finding supports the work of Vlasuk and co-workers who demonstrated the importance of polar residues close to the lipobox of *E. coli* Lpp, for translocation [39]. In the DOLOP database of predicted bacterial lipoproteins https://www.mrc-lmb.cam.ac.uk/genomes/dolop/predicted/ab.shtml [11] the algorithm does not include the requirement for this polar residue and thus for many of the proteins in this database, this residue is absent. These proteins are unlikely to undergo maturation to become lipoproteins and thus the size of bacterial lipoproteomes could be over-estimated. We also demonstrated experimentally in strain L91543 that the substitution of L at position 15 with F affected processing and for class 4 strains, it appears that a substitution even further from the lipobox; hydrophobic A at position 11 with T also affects processing. More experimental work is required for other bacterial lipoproteins to fully determine the structural features, in particular, critical AAs, that are required for lipoprotein maturation and localisation. From this, new programs with greater prediction power can be deployed which would be of importance to scientists developing vaccines based on bacterial lipoproteins, a growing area of research. Finally, our elucidation of the predominant vaccine target of Trumenba, and one of the targets of Bexsero, that is, unprocessed FHbp provides an important insight enabling a more intuitive interpretation of efficacy data.

## Methods

### Bacterial strains and culture conditions

*E. coli* strain JM109 single use competent cells were purchased from Promega and used for transformations in *E. coli*. BTH101 (Euromedex) was used as a reporter strain for BACTH assays. The 2 reference (control) strains of *N. meningitidis* used in this study are MC58, serogroup B:15:P1.7,16, ST-74; ET-5 purchased from LGC Standards and L91543 serogroup C:2aP1.2, ST-11; ET-37 kindly provided by Professor McFadden (University of Surrey). Strain H44/76, serogroup B:P1.7,16:F3-3: ST-32 was gifted by Rob Read and all other group B isolates were given by Christopher Bayliss with approval from Ray Borrow (Public Health England) (Table 2) and are listed in the Meningococcus Genome Library database (MRF collection). These isolates were obtained from patients in England, Wales, Northern Ireland and Scotland from 2009-2017.

*E. coli* strain strains were grown at 37°C in Lysogeny broth (LB) with shaking at 200 rpm or on agar (Merck). All meningococcal strains were grown on GC agar plates or GC broth (Difco) containing Kellogg’s glucose and iron supplements [49] in a moist atmosphere containing 5% CO2 at 37°C or at 30°C for transformation experiments and with shaking at 220 rpm for broth cultures. Antibiotics were purchased from Sigma and added at the following concentrations: kanamycin, 30 μg/ml and 60 μg/ml, erythromycin, 300 μg/ml and 0.3 μg/ml for *E coli* and *N. meningitidis* respectively; 100 μg/ml ampicillin for *E. coli;* and 30 μg/ml nalidixic acid for *E. coli* BTH101.

### Antibodies

Mouse anti-FHbp-Mabs, JAR4 and JAR5, were obtained from NIBSC. JAR4 is IgG2a and JAR5 is IgG2b; both were obtained from mice immunized with recombinant FHbp derived from MC58 [35]. Mouse anti-FHbp polyclonal antibody was kindly provided by Christoph Tang and rabbit anti-RecA IgG was purchased from Abcam. Factor H protein was purchased from Biorad and the mouse monoclonal IgG1 Factor H Antibody (OX24) conjugated with PE purchased from Santa Cruz Biotechnology. Secondary antibodies included donkey anti-rabbit HRP-linked IgG, sheep anti-mouse HRP-linked IgG (GE Healthcare, UK) for Western immunoblotting and rat anti-mouse IgG H+L conjugated with FITC for FACs analysis (Thermo Fisher Scientific).

### SDS-PAGE, Western immunoblotting and immuno-dot blotting

Whole cell lysates (WC) were prepared from broth or plate cultures resuspended in PBS fractionated by 12% or 16% (w/v) SDS-PAGE and immunoblotted as described previously [9]. For immunoblotting, membranes were incubated with JAR4 (diluted 1:1000 from 1 mg/ml stocks) or rabbit anti-RecA antibody diluted (1:1000) for 1 day, washed then incubated with either sheep anti-mouse or donkey anti-rabbit HRP-linked secondary antibody. Protein bands were detected by enhanced chemiluminescence (GE Healthcare, UK) or 3,3’,5,5’-tetramethylbenzidine (TMB) (Sigma). Band intensity was quantified using a GS-800™ calibrated densitometer (Bio-Rad) or ImageJ 1.x [50] calibrated to perform Optimal Density (OD) based on a pallet of colours in greyscale. Immuno-dot blots were performed as described previously [9].

### Palmitate Labelling of Lipidated Proteins

Bacterial cultures were grown in supplemented GC broth then after an initial doubling period, alkyne-labelled palmitic acid (Cayman Chemical) was added to a final concentration of 45 μM. Bacteria were incubated for at least 4 more hours at 37°C.

### Immuno-precipitation of FHbp from precipitated supernatant

Samples were immuno-precipitated with Protein G Mag Sepharose (GE Healthcare Life Sciences) and Mab JAR4. Following incubation of 100 µl of precipitated sample with 5 µg of JAR4 overnight at 4°C, samples were incubated for 1h with 100 µl of magnetic bead slurry. Using a Magnetic Particle Concentrator (MPC), beads were washed twice with PBS and FHbp recovered following the addition of 100 µl 0.1 M glycine-HCl (pH 2.5 to 3.1). Buffer exchange from glycine-HCl buffer to click reaction buffer (100 mM Na-Phosphate Buffer, pH 7) was performed with Slide-A-Lyzer Dialysis cassettes (Sigma).

### Click chemistry

The Click Chemistry labelling system “CuAAC Biomolecule Reaction Buffer Kit (THPTA based)” (Jena Bioscience) was used following manufacturer’s instructions [51]. FHbp was coupled to biotin azide (Stratech) then samples fractionated on 10-20% (w/v) SDS-PAGE gels (Novex). Western immunoblotting was performed by incubating the membrane with Streptavidin HRP-linked protein in PBS-BSA buffer and developing with TMB (Sigma).

### Harvesting cellular compartments of *N. meningitidis*

Periplasmic extracts were prepared using a method previously described [52]. Cells from over-night GC plate cultures were suspended to *A*600 1.0 in 500 µl buffer (50 mM Tris-HCl pH 8.0) and pelleted at 3,500 × g for 2 min. The pellet was resuspended in 200 µl of the same buffer and 20 µl of chloroform was added. After brief vortexing, tubes were incubated for 15 minutes at room temperature. After centrifugation at 6,500 × g for 2 min, the upper portion of the supernatant, containing the periplasmic proteins, was carefully aspirated and placed in a second tube. The pellet containing the remaining IM, OM and cytoplasmic proteins was partitioned following approaches adapted from 2 different research groups [53, 54]. The pellet was re-suspended in 500 µl buffer (50 mM Tris-HCl pH 8.0, 20% (w/v) sucrose) containing 1mg/ml lysozyme and incubated for 30 minutes at 4°C. After 2 cycles of freeze-thawing, the cells were subjected to sonication (2 bursts of 30s). Cellular debris were removed by centrifugation at 9,500 x g for 10 min. Ultra-centrifugation at 100,000 x g for 60 minutes enabled partitioning of the membranes in the pellet from cytoplasmic proteins in the supernatant. The IM was selectively solubilized by treatment with 200 µl of 1% (w/v) sodium lauroyl sarcosinate in 10 mM HEPES (N-2-hydroxyethylpiperazine N’-2-ethanesulfonic acid) pH 7.4 buffer. After centrifugation at 100,000 x g for 1 hour the supernatant containing solubilized IM proteins was separated from the pellet containing OM proteins. The pellet was washed with ethanol and resuspended in 200 µl PBS.

### Molecular Methods for DNA manipulations

Genomic DNA was extracted from *N. meningitidis* using the Gentra Puregene Yeast/Bact Kit (Qiagen) and plasmid DNA was extracted from *E. coli* using the QiaPrep Spin kit (Qiagen). DNA samples were analysed by agarose gel electrophoresis and visualised by staining with SYBR Safe (Invitrogen). Restriction enzymes (NEB), T4 DNA ligase (Promega), and Antarctic Phosphatase (NEB) were used according to the manufacturer’s recommendations. PCRs were performed using Q5 polymerase kit (NEB) in a Perkin- MJ Research PTC-200 Peltier Thermal Cycler or C1000 Touch™ Thermal Cycler (BioRad). Primers were purchased from Sigma and their sequences listed in Table 3. PCR products and restriction digested DNA were purified using the PCR Mini Elute kit (Qiagen). *E. coli* was transformed by heat shock [55].

**Table 3.**
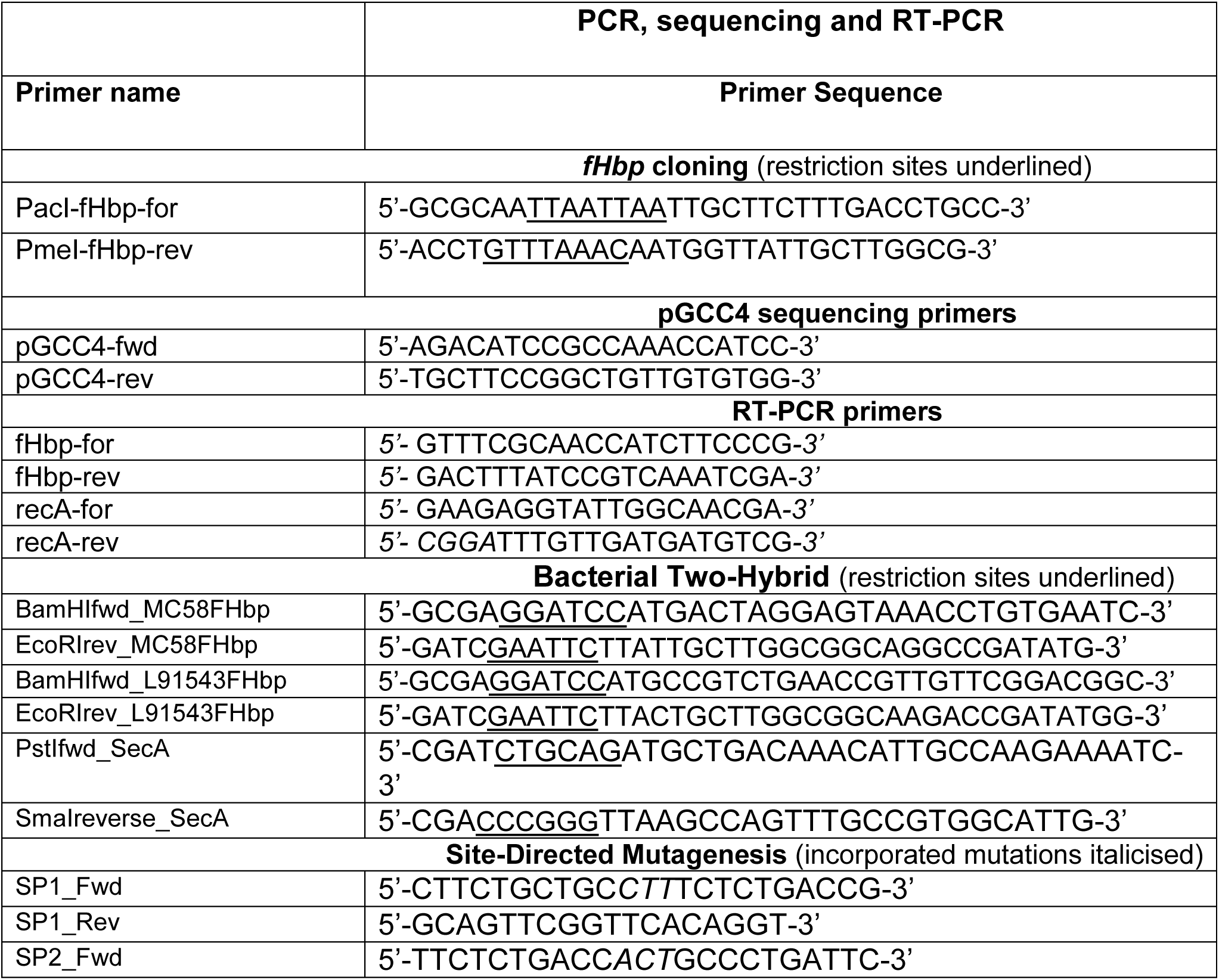
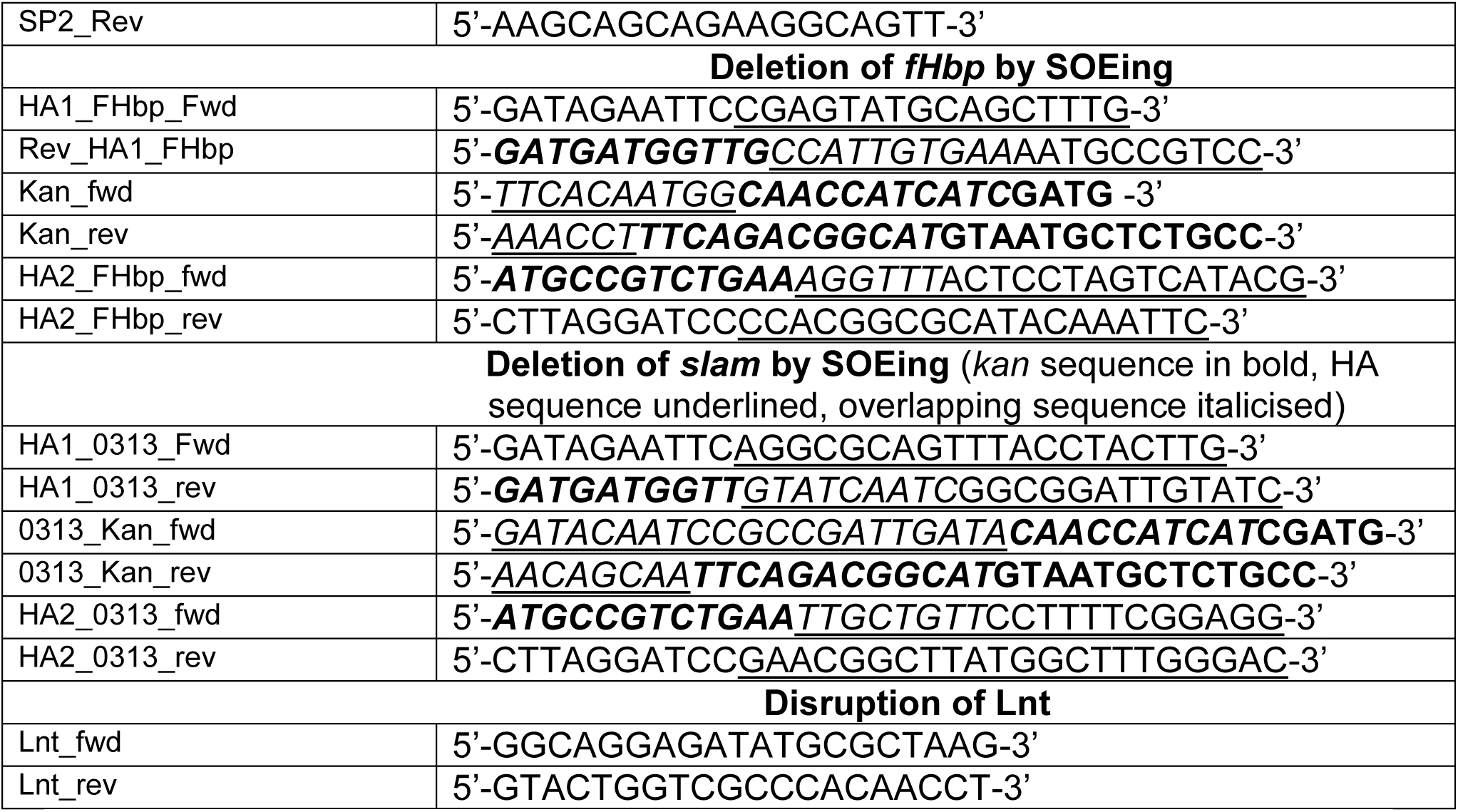
PCR primer pairs used in this study.

### Genome sequencing procedure

The *N. meningitidis* L91543 genome sequence was completed using a hybrid approach employing a combination of short and long reads assemblies produced by IonTorrent [28] and PacBio technologies. PacBio sequencing was conducted at TGAC (The Genome Analysis Centre, Norwich, UK) using an RSII sequencing machine with P6/C4 sequencing chemistry and a single SMRT cell. When analysing the PacBio assembly, long redundant regions were detected at the ends of the sequence. CLC Genomics Workbench software and high quality reads generated by IonTorrent PGM were used for elimination of artefactual redundancies and for circularisation, as well as for final verification of the sequence and correction of any errors produced by PacBio.

### BLAST analysis

Whole genome sequence (WGS) data from capsular group B isolates with complete FHbp profiles in the Meningitis Research Foundation Meningococcus Genome Library database (containing WGS data from all UK disease isolates from 2010-17) was interrogated using the BLAST tool implemented within the PubMLST.org/neisseria database using default settings (last analysed Sep 2018).

### Construction of strain L91543*fHbp*MC58

The region incorporating the promoter and open reading frame of *fHbp* was PCR-amplified from MC58 genomic DNA with primers PacI-*fHbp*-for and PmeI*-fHbp*-rev (Table 3) digested with PacI and PmeI and cloned into the PacI, PmeI sites of *Neisseria* complementation vector, pGCC4 (Addgene) [36]. The resulting plasmid, pGCC4*fHbp*MC58 was verified by DNA sequencing with pGCC4 primers (Table 3) and the plasmid used to transform *N. meningitidis* strain L91543 as previously described [56] with selection on erythromycin. A transformant was verified by PCR with subsequent DNA sequencing and designated L91543*fHbp*MC58. The same approach was used to create plasmid pGCC4*fHbp*L91543 with the L91543 version of *fHbp*, which was used for site-directed mutagenesis.

### Construction of strain L91543Δ*fHbp*

Gene Splicing by Overlap Extension (gene SOEing) was used to create a fusion PCR product to replace *fHbp* in strain L91543 with the kanamycin resistance gene (*kan*) from EZ::Tn5 <KAN-2> insertion kit (Epicentre) following the approach described by Horton [57]. In the first round of PCRs, homology arms (HA1 and HA2) of approximately 600 bp flanking *fHbp* were amplified from genomic DNA of L91543 using primers, HA1_FHbp_Fwd and Rev_HA1_FHbp and HA2_FHbp_fwd and HA2_FHbp_rev, and the *kan* gene was amplified using primers Kan_fwd and Kan_Rev. As shown in Table 3, in bold are the regions that bind to *kan*, underlined are the regions that bind to the HA of interest and in italics are the regions that overlap. The HA1 and *kan* PCR products obtained were gene-cleaned then used as template for the second round of PCR with primers HA1_FHbp_Fwd and Kan_Rev and the annealed product generated cleaned and used as template along with the HA2 PCR product for a third round of PCR with primers HA1_FHbp_Fwd and HA2_FHbp_rev. The final PCR product generated containing HA1-*kan*-HA2 was gene cleaned and sequenced for confirmation. The verified construct was used to transform strain L91543 with selection on kanamycin. Deletion mutants were confirmed by PCR and DNA sequencing and designated L91543Δ*fHbp*.

### Disruption of Slam

In the same manner as that described above, the gene encoding Slam (NMB0313) was replaced with *kan*. For the first round of PCRs, primers HA1_0313_Fwd and HA1_0313_rev and HA2_0313_fwd and HA2_0313_rev were used to amplify approximately 600 bp of sequence flanking *slam* from genomic DNA of MC58 and *kan* was amplified with primers 0313_Kan_fwd and 0313_Kan_rev (Table 3). The appropriate primer pairs were used in second and third round PCR and the construct confirmed by sequencing then used to transform MC58, L91543 and isolates 6 and 18 with selection on kanamycin. Insertion mutants were confirmed by PCR and DNA sequencing and designated MC58Slam, L91543Slam, Isolate6Slam, and Isolate18Slam.

### Disruption of Lnt

Primers, Lnt_fwd and Lnt_rev (Table 3), annealing approximately 600 bp upstream and downstream respectively of *lnt* in strain MC58Lnt, were used to amplify *lnt* that had been disrupted by the insertion of Tn5<KAN-2> (Epicentre) [6]. The PCR product generated was used to transform L91543 and isolates 6 and 8 with selection on kanamycin. Deletion mutants were confirmed by PCR and DNA sequencing and designated L91543Lnt, Isolate6Lnt and Isolate18Lnt.

### Site-Directed Mutagenesis

Site-directed mutagenesis of the *fHbp* SP in pGCC4*fHbp*L91543 to repair SNP1 and SNP2 individually was performed using the Q5® Site-Directed Mutagenesis Kit (NEB) according to the manufacturer’s recommendations. SNP1 was corrected to create plasmid pGCC4L*fHbp*SNP1 and SNP2 was corrected to create plasmid pGCC4L*fHbp*SNP2. Seven ng of pGCC4*fHbp*L91543 and 0.5 µM of each pair of primers (Sigma) were used for amplification and incorporation of the desired mutation. Primers SP1_Fwd and SP1_Rev, and SP2_Fwd and SP2 _Rev (Table 3) were used to incorporate SNP1 and SNP2 respectively. PCR was performed in a C1000 Touch™ Thermal Cycler (BioRad) with the following thermo-cycling conditions; 98°C for 30 s followed by 25 cycles of 98°C for 10 s (denaturation), 68°C for 15 s (annealing), 72°C for 3 minutes and 25 s (extension) and 72°C for 2 minutes (final extension). One μl of the PCR product was kinase-, ligase- and DpnI-treated (KLD treatment) and incubated for 5 minutes at room temperature. Five μl of the KLD mix was then used to transform competent cells. After plating on LB agar with kanamycin and incubating overnight, several colonies were isolated, grown individually in LB broth with kanamycin and plasmid DNA extracted and sequenced with pGCC4 primers (Table 3) for verification. To repair both SNPs, the PacI-PmeI fragment of pGCC4*fHbp*L91543 was commercially synthesized (Life Technologies) with the 2 SNPs repaired to resemble the *fHbp* SP of MC58 and the DNA cloned into the PacI-PmeI sites of pGCC4 to create pGCC4L*fHbp*SNP1+2. The construct was verified by sequencing with pGCC4 primers (Table 3).

Strain L91543Δ*fHbp* was transformed with pGCC4*fHbp*L91543 to create L91543Δ*fHbp*L*fHbp* with no SNP corrections as a negative control. The same strain was transformed with pGCC4L*fHbp*SNP1 to generate recombinant strain L91543Δ*fHbp*+L*fHbp*SNP1, with pGCC4L*fHbp*SNP2 to generate strain L91543Δ*fHbp*+L*fHbp*SNP2 and finally with pGCC4L*fHbp*SNP1+2 to create strain L91543Δ*fHbp*+L*fHbp*SNP1+2.

### Bacterial Two-Hybrid Assay (BACTH)

The protein–protein interaction of MC58 FHbp and L91543 FHbp with SecA was investigated using the Bacterial Adenylate Cyclase Two-Hybrid System Kit (Euromedex) according to manufacturer’s instructions. First, *fHbp* from MC58 and from L91543 was PCR-amplified with the primer pair BamHIfwd_MC58FHbp and EcoRIrev_MC58FHbp and primer pair BamHIfwd_L91543FHbp and EcoRIrev_L91543FHbp respectively (Table 3) then the PCR products cloned separately into vector pUT18. The gene encoding SecA from MC58 was PCR-amplified with primer pair, PstIfwd_SecA and SmaIreverse_SecA (Table 3) then cloned into vector pKT25. 25-50 ng of the appropriate prey (pKT25-based construct) and the equivalent concentration of appropriate bait (pUT18-based construct) were co-transformed into 100 μl of competent *E. coli* BTH101 cells and plated on MacConkey agar containing 0.5 mM IPTG and appropriate antibiotics. Bacteria expressing interacting hybrid proteins formed pink/purple colonies while cells expressing non-interacting proteins remained white/light pink. As a positive control, a co-transformant containing commercial pKT25-zip and pUT18-zip constructs was used. Co-transformants containing empty vector pKT25 and/or pUT18 were used as negative controls. Pink colonies were isolated and grown in LB broth and plasmid DNA extracted for verification by PCR and sequencing.

### β-Galactosidase Assay

Following the approach previously described [58], to measure the level of protein-protein interaction between FHbp from MC58 or L91543 with SecA, LB broth cultures were grown to *A*600 0.6 then induced for 3 hours with 0.5 mM IPTG. Five μl of induced culture were mixed with 900 μl of Z buffer (0.06 M Na2HPO4 2H2O, 0.04 M NaH2PO4, 0.01 M KCl, 2 mM MgSO4.7H2O, 14.20 M β-mercaptoethanol (pH 7), before addition of 20 μl of 0.1% (w/v) SDS and 100 μl of CHCl3 to permeabilise the cells. Substrate solution was prepared by solubilising orthonitrophenyl-β-galactosidase (Sigma) in Z buffer without the β-mercaptoethanol to a final concentration of 4 mg/ml. 40 μl of substrate solution were added to 180 μl of Z buffer and 20 μl of permeabilised cells in a 96-well plate. The plate was incubated at room temperature for at least 20 minutes. Readings were taken on a microplate reader at *A*405 and *A*540 and β-galactosidase activity calculated using the equation below and expressed in Miller units.

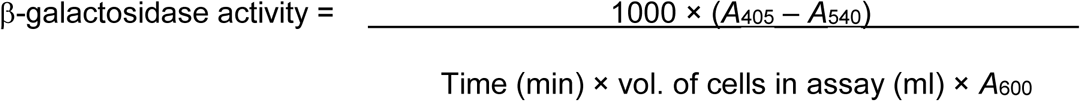

### FACS analysis of isolates to measure the number of JAR4 antibodies bound per cell (ABC) and quantify surface localisation of FHbp

FHbp surface expression was assessed using a MoFlo Astrios EQ, Cell Sorter (Beckman Coulter). The approach used was similar to that described previously [23]. Approximately 1 × 10^5^ bacteria were suspended in PBS containing 1% (w/v) bovine serum albumin (PBS-BSA) and incubated with JAR4 in a final volume of 100 μl for 1 hour at 37°C. After two washes with PBS, JAR4 binding was detected using FITC-conjugated rat anti-mouse IgG (H+L) (Thermo Fisher) at a 1:50 dilution for an incubation period of 1 h. After the final 2 washes with PBS, samples were resuspended in 500 μl of PBS-BSA with 4% (w/v) formalin. The negative control consisted of cells incubated with the secondary antibody alone. Quantum™ Simply Cellular® anti-rat (Bang Laboratories) microspheres were used to determine the number of ABC according to the manufacturer’s instructions. Median channel values for each population of microspheres were acquired for entry into the QuickCal® spreadsheet to generate a curve that enabled the acquisition of ABC for each isolate.

### FACS analysis of isolates with anti-fH antibody to quantify fH binding

Approximately 1 × 10^5^ bacterial cells were suspended in PBS containing 1% (w/v) PBS-BSA and incubated with fH (Biorad) at a final concentration of 5 μg/mL for 1 hour at room temperature. After two washes with PBS, anti-fH antibody (OX-24) conjugated to PE was added at a 1:50 dilution for 1 h. After 2 more washes in PBS, samples were resuspended in 200 μl of PBS-BSA with 4% (w/v) formalin. The negative control consisted of cells incubated with anti-fH antibody alone. The relative Mean of Fluorescence Intensity (rMFI) for total fluorescent cells was acquired for each sample and these values normalised against rMFI of cells incubated with secondary antibody alone.

### Susceptibility of isolates to killing in Serum Bactericidal Antibody assays

The susceptibility of the isolates to complement-mediated killing with antibodies JAR4 and JAR5 was assessed using approaches adapted from previous studies [42, 59, 60]. JAR4 and JAR5 were used in combination, in place of serum antibodies, to generate bactericidal activity [41–43]. Eight μg of each antibody were used to ensure maximum killing [42]. Complement was obtained from lyophilized human sera (Sigma) reconstituted in sterile water as per manufacturer’s instructions. Complement and antibodies were added to IgG-depleted human sera (Stratech) that had been heat-inactivated for 30 minutes at 56°C. The bactericidal activity i.e. the percentage of killing by the antibodies for each isolate was then determined from the CFU counts after 60 minutes incubation in the reaction mixture and compared with CFU counts from negative control wells at time zero.

### Affect of globomycin treatment on FHbp

Following a similar approach to that previously described [18] GC broth un-supplemented with iron was inoculated with meningococci cells from freshly grown plate cultures and cells incubated until densities reached *A*600 0.1. The cells were then incubated with 10 µg/ml of globomycin (Sigma) for 2 h. Suspensions were adjusted to *A*600 0.4 then cells harvested and lysed in SDS-sample buffer before fractionation by SDS-PAGE and immunoblotting with JAR4.

### Reverse Transcription PCR (RT-PCR)

Following a similar approach to that we described previously [9], RNA was extracted from 1 ml cell suspensions of each isolate standardised to *A*600 0.65 (containing approximately 2 x 10^8^ cells) using the RNeasy Mini kit (Qiagen) with enzymatic lysis and Proteinase K digestion. On-column DNA digestion was performed using the RNase Free DNase kit (Qiagen). RNA quality was assessed for genomic contamination and integrity using a NanoDrop Lite (Thermo Fisher Scientific) and running 1 µl of sample on a 1% agarose gel.

One µg of cDNA was synthesised using the QuantiTect reverse transcription kit (Qiagen) with the initial genomic wipe-out step included. RT-PCR was performed in a 50 µl reaction mixture with One Taq 2x master mix (NEB), *2*0 ng of cDNA and 0.2 µM of each primer (Sigma). For amplification of cDNA of *fHbp*, *fHbp*-for and *fHbp*-rev primers were used and for amplification of *recA*, *recA*-for and *recA*-rev primers were used (Table 3). To qualitatively check for different levels of expression, PCR was performed in a C1000 Touch™ Thermal Cycler (BioRad) with the following thermo-cycling conditions; 94°C for 30 s followed by 35 cycles of 94 °C for 15 s (denaturation), 56°C for 15 s (annealing) and 68°C for 20 s (extension). PCR products were visualised on a 1% agarose gel.

### Statistics

Data are shown as mean ± SEM. Multiple comparisons between groups were performed by one way ANOVA followed by Dunnett’s test. Unpaired T tests with Welsh’s corrections were used for comparisons between 2 groups. A value of P ≤ 0.05 was considered statistically significant. Post hoc tests were run if F achieved P < 0.05 and there was no significant variance. All statistical analyses were performed in GraphPad Prism version 6.00 for Windows. No statistical methods were used to predetermine sample size. The experiments were not randomized other than for selection of clinical isolates used in this study. The investigators were not blinded to allocation during experiments and outcome assessment.

## Data availability

The FACS data discussed in this publication have been deposited in FlowRepository [61] and are temporarily available through accession numbers;

FR-FCM-ZYTU,

http://flowrepository.org/id/RvFrLUcAM69ZrTziXZJn622NsvQgwRuVqoRC28XA9wBxR2KZsAJ2yym5ToAfMt6p,

FR-FCM-ZYTV,

http://flowrepository.org/id/RvFryhGbJHv5UMqqZ1bEfLEz6aO8QEV0EsQtjKdcSjlMt1rCtFIiT9HmOrMZYBms and

FR-FCM-ZYUZ

http://flowrepository.org/id/RvFraaVCkdcJJYpYJOc4CFQNbETeVGwTy1bcNwXpA3YotemW9trGUDNKJFgxXnKi.

The closed genome sequence of L91543 is available under accession number CP016684.

## Acknowledgements

We thank Conselho Nacional de Desenvolvimento Científico e Tecnológico (CNPq) for funding RAG da Silva’s work on this project (201521/2015-6). This publication made use of the MRF Meningococcus Genome Library (http://www.meningitis.org/research/genome) developed by Public Health England, the Wellcome Trust Sanger Institute and the University of Oxford as a collaboration. We thank Professor Nigel P Minton for bench space and use of equipment, Professor Liz Sockett for proof-reading this manuscript, Izadora L. A. Rabelo for assisting with FACS analysis and Marcelo Depólo for helpful discussions about SPs.

## Author contributions

RAGS performed all of the experiments, analysed the data and contributed to the experimental design and writing of the manuscript, RG devised the project, supervised the experiments, analysed the data and wrote the manuscript, AVK closed the genome of L91543, NJO and KGW assisted with the Bioinformatics analyses, CDB provided the strains and FHbp protein sequences and AR provided technical advice.

---

**Figure S1.**
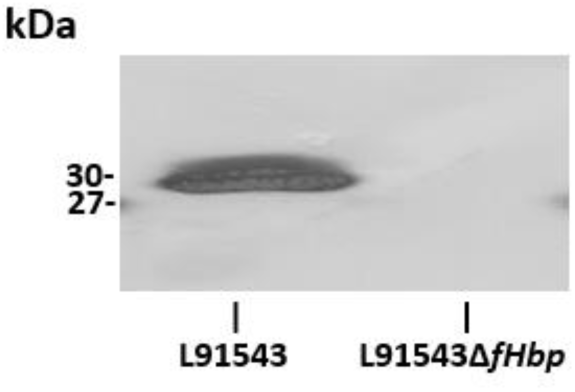
Western immunoblotting of WCL of L91543 and L91543Δ*fHbp* with JAR4.

**Table S1.**
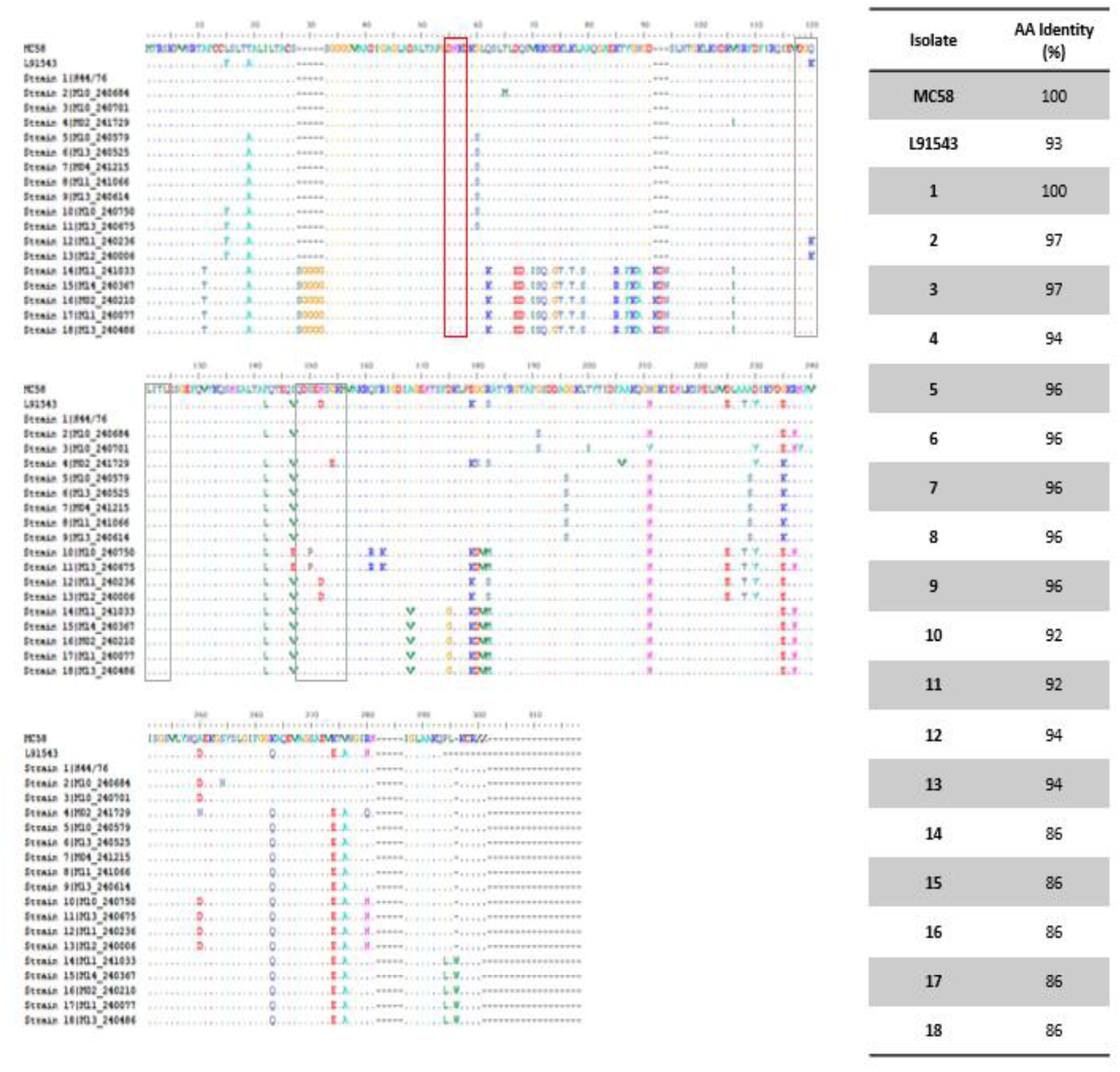
Multiple alignment of FHbp of isolates used in this study and amino acid identities across whole protein relative to MC58. Framed in red and grey boxes are AA residues in the epitopes recognised by JAR4 and JAR5 respectively.

**Table S2.**
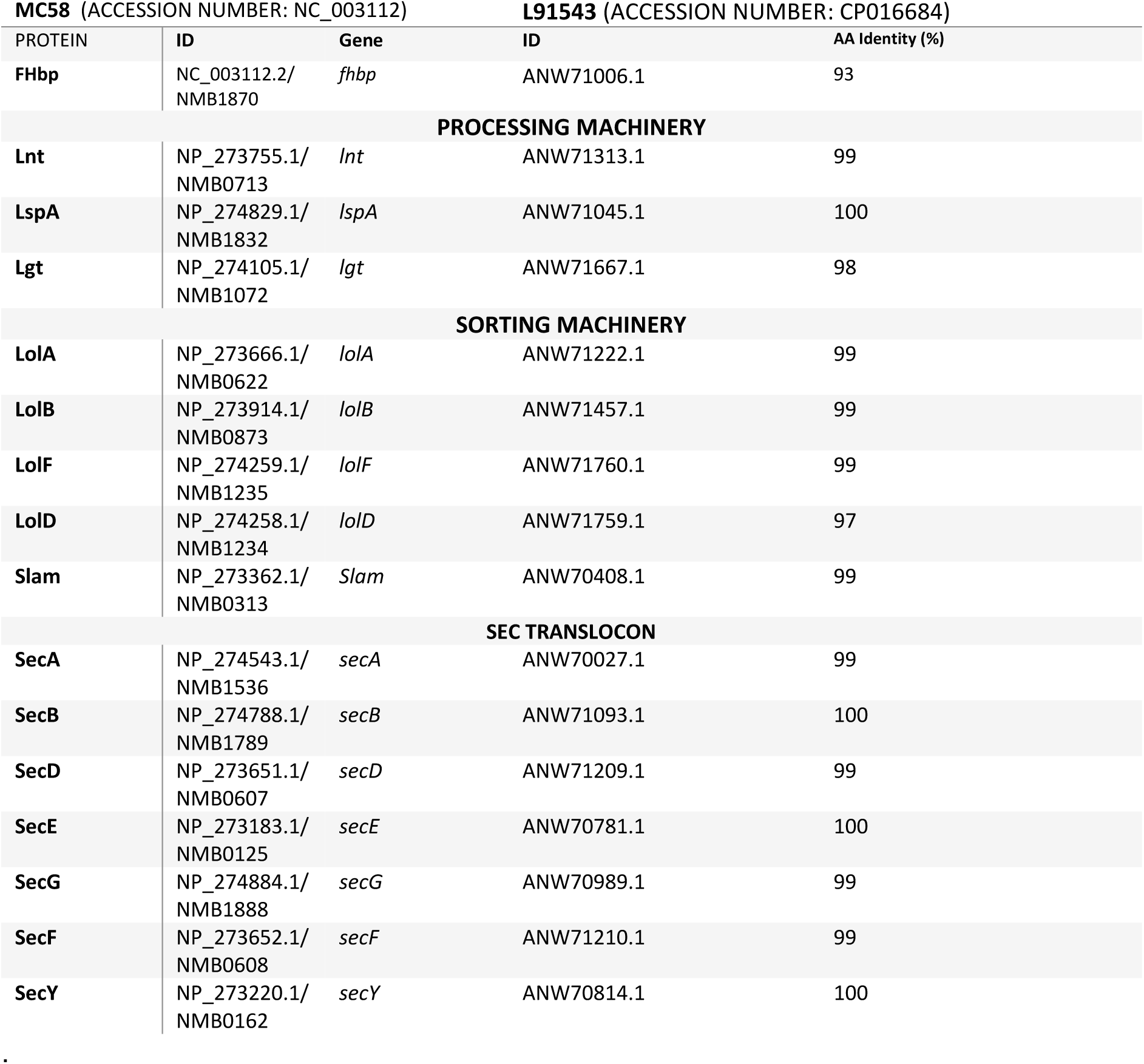
Amino acid identity of proteins involved in translocation, processing and surface localisation of FHbp in L91543 compared to MC58 [9]

## Notes

http://flowrepository.org/id/RvFrLUcAM69ZrTziXZJn622NsvQgwRuVqoRC28XA9wBxR2KZsAJ2yym5ToAfMt6p

http://flowrepository.org/id/RvFryhGbJHv5UMqqZ1bEfLEz6aO8QEV0EsQtjKdcSjlMt1rCtFIiT9HmOrMZYBms

http://flowrepository.org/id/RvFraaVCkdcJJYpYJOc4CFQNbETeVGwTy1bcNwXpA3YotemW9trGUDNKJFgxXnKi

